# Coronavirus endoribonuclease nsp15 induces host cellular protein synthesis shutoff

**DOI:** 10.1101/2023.03.20.533404

**Authors:** Xiaoqian Gong, Shanhuan Feng, Bo Gao, Shouguo Fang, Wenlian Weng, Wenxiang Xue, Hongyan Chu, Yanmei Yuan, Yuqiang Cheng, Yingjie Sun, Lei Tan, Cuiping Song, Xusheng Qiu, Chan Ding, Min Liao, Edwin Tijhaar, Maria Forlenza, Ying Liao

## Abstract

The endoribonuclease (EndoU) nsp15 of coronaviruses plays an important role in evasion of host innate immune responses by reducing the abundance of viral double-stranded RNA, whereas less is known about potential host cellular targets of nsp15. In this study, we show that cellular protein synthesis is inhibited upon over-expression of nsp15 from four genera of coronaviruses and this is accompanied by nuclear retention of the poly(A) binding protein cytoplasmic 1 (PABPC1). We also show that the EndoU activity of nsp15 is indispensable for both, inhibition of protein synthesis and PABPC1 nuclear relocation. FISH analysis using oligo-dT probes, revealed an overlap between the localization of cellular mRNA and that of overexpressed nsp15 in some cells, suggesting that, when expressed alone, nsp15 may target host mRNA. When investigating the association of nsp15 on protein shut off in the context of a viral infection, we observed that the *γ*-coronavirus infectious bronchitis virus (IBV), induced host translation shutoff in an p-eIF2α-independent manner and mainly retained PABPC1 in the cytoplasm, whereas the nsp15 EndoU-deficient IBV accumulated viral dsRNA and caused p-PKR-p-eIF2α-dependent host protein translation shutoff, accompanied with PABPC1 nuclear relocation or stress granule (SG) localization. This phenomenon suggests that during infection with wild type IBV, although the cellular translation is inhibited, initiation of viral mRNA translation leads to PABPC1 binding to viral mRNA, thereby preventing its nuclear entry; during infection with nsp15 EndoU-deficient IBV however, the eIF2α-dependent host protein translation shutoff prevents both host and viral mRNA translation initiation, releasing PABPC1 from binding to cytosolic and viral mRNA, thereby relocating it to the nucleus or to SG. Altogether, this study reveals unique yet conserved mechanisms of host protein shutoff that add to our understanding of how coronaviruses regulate host protein expression through a mechanism that involves catalytically active nsp15 EndoU, and describes how nsp15 may target both, viral and host mRNA.

**Author summary:** It has been reported that coronavirus infection suppresses host protein translation, *α-* and *β-* coronavirus nsp1 is responsible for inhibition of host gene expression. However, for *γ-* and *δ*-coronavirus, there is no nsp1 and the underlying mechanisms by which virus regulates host translation are not well characterized. Here, we show that coronavirus endoribonuclease nsp15 is responsible for the inhibition of host translation by targeting to host factors, meanwhile it helps virus bypass the PKR-eIF2α mediated host translation shutoff, which is harmful for virus gene expression, by reducing the accumulation of viral dsRNA. This novel finding gives insight how does nsp15 targets to both host factors and viral RNA, to facilitate virus replication. Moreover, the novel function of nsp15 is found to be conserved among coronaviruses, revealing the essential role of this endoribonuclease in hijacking host translation machinery for virus replication.

## Introduction

Coronaviruses are classified into four genera, *α, β, γ* and *δ*. They are enveloped, positive-sense, single-stranded RNA viruses possessing the largest known RNA genome (approximately 25 to 32 kilobases) [1]. Over two-thirds of the genome at the 5’-end comprises the open reading frames ORF1a and ORF1b; the translation of ORF1b requires a programmed −1 ribosomal frameshifting mechanism [2]. Upon entry into host cells, coronavirus genome serves as the template for the translation of polyproteins 1a and 1ab, which are cleaved by internal papain-like protease (nsp3) and 3C-like protease (nsp5), to produce non-structural proteins (nsp) [3]. The *α* and *β* coronaviruses encode 16 nsps (nsp1 to nsp16) [4]; while the *γ and δ coronaviruses* only encode 15 nsps (nsp2-nsp16) and lack the most N-terminal cleavage product nsp1 [5, 6]. A number of nsps contain domains involved in transcription and replication of viral RNA, including the RNA-dependent RNA polymerase (RdRp) nsp12, the template-primer prividers nsp7 and nsp8, the RNA helicase/5’-triphosphatase nsp13, the exoribonuclease nsp14, the endoribonuclease (EndoU) nsp15, and the RNA-cap methyltransferase nsp14 and nsp16 [7–13]. Amongst them, RdRp nsp12 plays a central role to drive the replication and transcription of viral RNA, while the other nsps play a supportive role.

It has been reported that the highly conserved EndoU nsp15 is an integral component of the replication and transcription complex (RTC) [14–16], where it possesses uridylate-specific endonucleolytic activity on viral RNA. The RTC is accommodated in virus-induced double membrane vesicles (DMV) during SARS-CoV-2, SARS-CoV, MERS-CoV or MHV infection [17–21], or in zippered ER and spherules single membranes during IBV infection [22], which is formed by the transmembrane protein nsp3, nsp4, and nsp6 [23, 24]. The viral ligand double stranded (ds)RNA, an intermediate product of viral replication, is as well present in the virus-induced remodelled intracellular membrane structures [19, 25]. Nsp15 activity contributes to keeping the amount of dsRNA low to evade the detection by host cell sensors. Such role for nsp15 has been reported for MHV, HCoV-229E, PEDV, IBV [26–29], and likely also for SARS-CoV-2 [30, 31]. While the role of nsp15 in innate immune escape is well established, its effect on host cells remains poorly understood. In our previous study, we observed that nsp15 suppresses the chemically- or physically-induced formation of stress granules (SG), and promotes nuclear accumulation of the cytoplasmic poly(A) binding protein (PABPC1) [29]. This observation suggests that nsp15 not only targets viral RNA, but also regulates the host functions by targeting some unknown host substrates.

In order to successfully replicate, coronaviruses employ a range of strategies to escape or antagonize the host immune responses [32]. Inhibition of host gene expression is not only an alternative strategy to antagonize the host innate immune response by reducing the synthesis anti-viral proteins, but also a smart way to hijack the host translation machinery to facilitate the translation of viral mRNA instead. The process of eukaryotic gene expression includes transcription, RNA processing, nuclear export of RNA, protein translation, and post-translational modification [33]. Viruses may suppress the host gene expression by reducing the levels of host mRNA or preventing their association with ribosomes or translation initiation factors. For example, poliovirus 3C protease inhibits RNA polymerase II mediated transcription initiation by cleavage of transcription activator Oct-1 [34, 35]; influenza A virus (IAV) polymerase acidic (PA) protein snatches the capped primers from nascent host transcripts for the synthesis of viral mRNA [36]; IAV NS1 inhibits polyadenylation of cellular precursor mRNA (pre-mRNA) and prevents the nuclear export of cellular mRNA [37–39]; human immunodeficiency virus (HIV) viral protein R (VPR) inhibits the splicing of host pre-mRNA [40]; herpes simplex virus (HSV) infected cell protein 27 (ICP27) blocks host transcription termination [41]; poliovirus 2A protease affects cellular mRNA nuclear export by mediating nucleoporin cleavage [42]; poliovirus 3C proteinase cleaves PABP to inhibit translation initiation [43]. It has been reported that several *α-* and *β-*coronaviruses, through their nsp1, hijack the host translation machinery by repressing host mRNA transcription in the nucleus [44], preventing the nuclear export of host mRNA [45, 46], degrading host mRNA in the nucleus and cytoplasm [44, 47–50], and inhibiting host mRNA translation by interfering ribosome [49, 51, 52]. However, for the *γ-*coronavirus IBV and *δ -*coronavirus PDCoV, which lack nsp1, although host protein translation shutoff is also observed, the protein(s) potentially involved in are not well characterized [53–55], although the IBV 5b and S were identified to be associated with translation inhibition [56, 57].

Several viral endonucleases play a role in regulating not only viral RNA but also host gene expression by targeting host mRNA for degradation. For example, the *α-herpesvirinae* herpes simplex virus (HSV) virion host shutoff (Vhs) protein [58–60], the *γ-herpesvirinae* Kaposi’s sarcoma-associated herpesvirus (KSHV) shutoff and exonuclease (SOX protein) [61–63], Epstein Barr virus (EBV) BGLF5 protein [64–66], and murine herpesvirus 68 (MHV-68) muSOX protein [61, 67], possess endonuclease activity and inhibit synthesis of cellular proteins by promoting the global mRNAs degradation [61, 63, 68, 69]. As mentioned, the IAV cap-dependent endonuclease PA is associated with the RNA polymerase complex and is responsible for snatching capped oligonucleotides from cellular pre-mRNAs and for using them as primers for the synthesis of viral mRNAs, thereby helping viral gene expression while suppressing host mRNA maturation [36, 70, 71]; moreover, IAV encodes another endonuclease, PA-X, that selectively degrades host RNAs and usurps the host mRNA processing machinery to destroy nascent mRNAs and limit host gene expression [72–74].

Here we investigated the ability of EndoU nsp15 of IBV and of other coronaviruses to inhibit cellular protein synthesis. We provide evidences that nsp15 inhibits host protein translation through a mechanism that involves targeting host factors involved in the translation complex, including host mRNA and PABPC1; a mechanism for which the EndoU activity is indispensable. In the context of IBV infection, nsp15, probably in concert with other IBV proteins, trigger an eIF2α-independent host translation shutoff. When nsp15 EndoU activity is deficient, a higher amount of viral dsRNA accumulates in the cells [29], the nsp15 deficient IBV inhibits host protein translation through a PKR-eIF2α-dependent mechanism. Altogether, our findings unveil new, yet conserved strategies shared among coronaviruses, to regulate host gene and protein expression and the role of the EndoU nsp15 therein.

## Results

### IBV nsp15 inhibits exogenous protein synthesis but does not affect mRNA levels

We previously reported that IBV nsp15 interferes with the chemically- and physically-induced SGs formation, possibly by targeting the host translation machinery [29]. Furthermore, during a screening of IBV-encoded type I IFN antagonists using a luciferase reporter system, we observed that IBV nsp15 reduced the expression not only of the IFNβ promotor-driven luciferase, but also the expression of the co-transfected plasmids encoding HA-tagged MAVS (HA-huMAVS) in 293T cells (**supplementary Fig 1**). These data prompted us to hypothesize that IBV nsp15 interferes with the host translation system. It was noted that 5a and E also suppressed the co-transfected huMAVS expression, meanwhile reduced the expression of IFNβ-driven luciferase. This result indicates IBV encodes several proteins involved in inhibition of protein expression.

To examine whether IBV nsp15 indeed interferes with the host protein synthesis, DF-1 cells were co-transfected with plasmids encoding Flag-tagged IBV nsp15 and V5-tagged constitutively active form (N-terminal domain) of chicken MDA5 [V5-chMDA5(N)] [75], nsp15 and HA-chMAVS [76], nsp15 and V5-tagged chicken interferon regulatory factor 7 (V5-chIRF7) [77], or nsp15 and enhanced green fluorescent protein (EGFP). Vector PXJ40, Flag-tagged nsp7, nsp8, nsp9, and nsp13 were co-transfected with corresponding plasmids as control. Western blot analysis showed that overexpression of nsp15 strongly suppressed the protein levels of V5-chMDA5(N), HA-chMAVS, V5-chIRF7, and also EGFP to various degrees (**Fig 1A**), whereas overexpression of nsp7, nsp8, nsp9, nsp12, or transfection of PXJ40 did not. These results confirm that nsp15 inhibits the expression of exogenous proteins encoded by co-transfected plasmids.

**Fig. 1.**
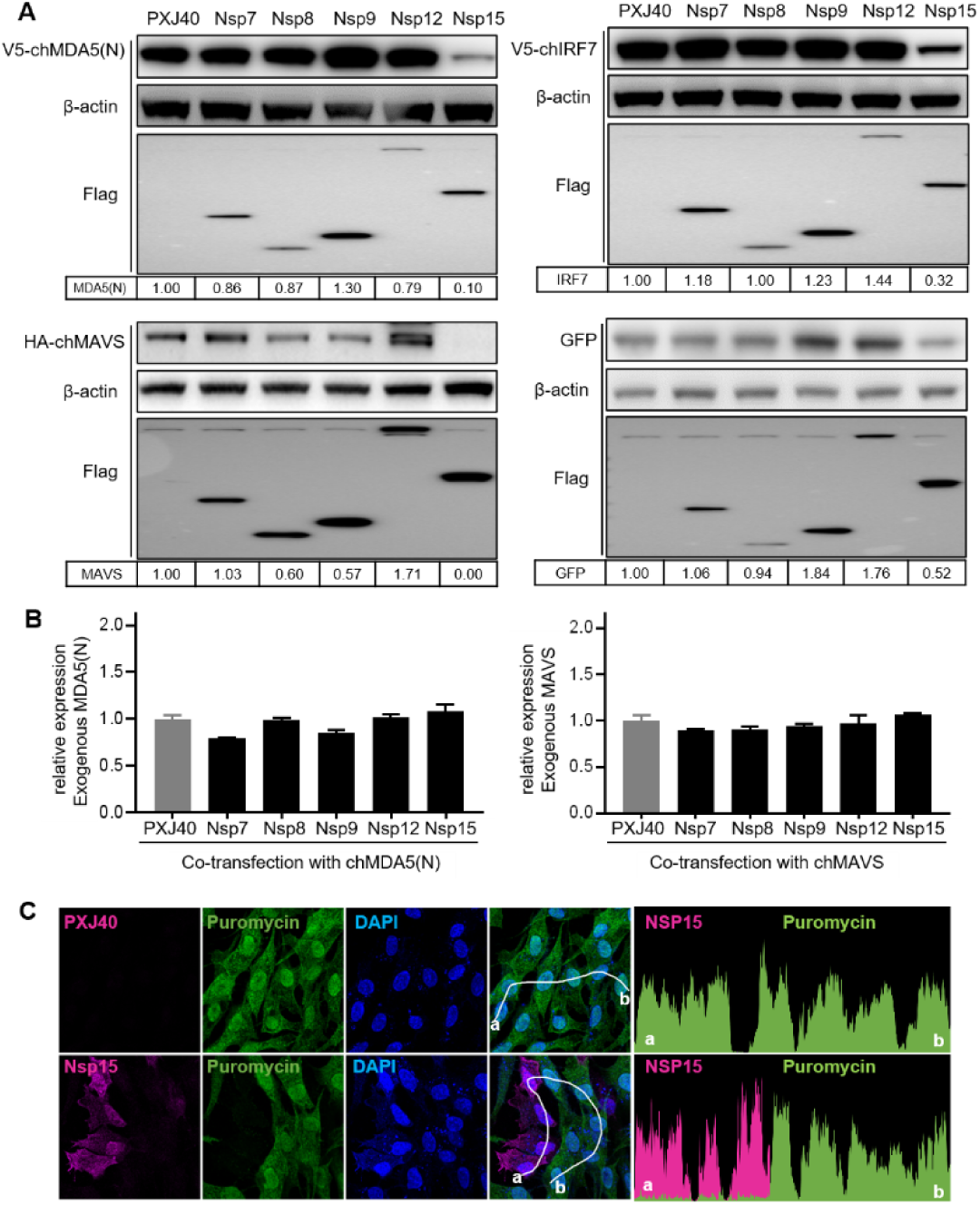
IBV nsp15 inhibits *de novo* protein synthesis but does not affect mRNA levels. (A) DF-1 cells were co-transfected with plasmids encoding Flag-tagged IBV nsp7, nsp8, nsp9, nsp12 or nsp15 and the plasmids encoding a constitutively active form of V5-chMDA5(N), or HA-chMAVS, or V5-chIRF7, or enhanced green fluorescent protein (EGFP). The empty PXJ40 vector was included as a control. After 24 h, cells were collected for western blot analysis. Protein signals were detected using the indicated antibodies, and β-actin was detected as loading control. The density of the protein bands was analysed with ImageJ, normalized by the density of β-actin, and the ratio was presented relative to the density detected in the corresponding PXJ40 sample. (B) PXJ40, or Flag-tagged IBV nsp7, nsp8, nsp9, nsp12 or nsp15 were co-transfected into DF-1 cells with V5-chMDA5(N) or HA-chMAVS. After 24 h, cells were collected and the RNA were extracted, subjected to quantitative RT-PCR, using primers spanning the tag (V5 or HA) sequence and the chMDA5(N) or chMAVS sequence. mRNA levels of V5-chMDA5(N) or HA-chMAVS were normalized relative to the β-actin housekeeping gene and presented relative to PXJ40 group. Values present results of one representative experiment, which was performed three times with comparable results. Error bars indicate standard deviation of triplicate values within one experiment. (C) DF-1 cells were transfected with plasmid encoding Flag-nsp15 or PXJ40 for 23 h and treated with puromycin (5 µg/ml) for 1 h to label *de novo* synthesized peptides. Indirect immunofluorescence was performed to detect nsp15 (magenta), puromycin (green), and nuclei (DAPI, blue). Fluorescence intensity of nsp15 and puromycin in individual cells along the white line (from a to b) is shown in the right panel (histogram plot).

Considering the EndoU activity, we next asked whether the effects of IBV nsp15 on exogenous protein expression were mediated by reducing mRNA levels. To this end, we quantified the transcripts derived from the co-transfected plasmids encoding V5-chMDA5(N) or HA-chMAVS. To prevent amplification of endogenous transcripts, primers were designed to span the V5 or HA tag sequence. Using random primers for an unbiased cDNA synthesis, the quantitative RT-PCR analysis showed that nsp15 did not affect mRNA levels of transcripts derived from co-transfected chMDA5(N) or chMAVS (**Fig 1B**). These data are also consistent with our previous report showing that during IBV infection no effect on host mRNA stability was observed [53].

Since the nsp15 induced inhibition of exogenous protein expression was not restricted to proteins involved in anti-viral responses but was also observed for EGFP, we next asked whether the effect on protein synthesis is universal to host endogenous proteins. To test this, we determined the signal of puromycin-labelled, *de novo* synthetized, endogenous peptides. Puromycin is an analogue of tRNA that binds to growing peptide chains and causes the release of premature peptide chains [78]. Therefore, the signal of puromycin-labelled peptides, detected with an anti-puromycin antibody, represents the number of peptides *de novo* synthesized during the period of puromycin treatment. In all nsp15-expressing cells, puromycin labelling signal was strongly reduced, while this was not observed in nsp15 non-expressing cells, or cells transfected with the control PXJ40, indicating that nsp15 indeed suppresses *de novo* protein synthesis (**Fig 1C**). Together, these results show that overexpression of nsp15 inhibits the global synthesis of endogenous proteins.

### The catalytic activity and oligomeric structure of IBV nsp15 are indispensable for inhibition of *de novo* protein synthesis

The two conserved catalytic residues for IBV nsp15’s EndoU activity reside at histidine residues H223 and H238, and the two residues critical for oligomerization resides at aspartic acid D285 and D315. We previously reported that also for IBV, the EndoU activity of nsp15 is required to limit the accumulation of dsRNA intermediates in the cells, thereby escaping host recognition and delaying IFNβ production [29]. To investigate whether the EndoU activity is required also for the observed effects on *de novo* protein synthesis inhibition, we made use of previously generated mutated nsp15 in which either one of the catalytic histidine (H) or one of the aspartic acid (D) residues, were substituted by an alanine (A) residue (H223A, H238A, D285A, D315A) [29]. Western blot analysis revealed that mutating the catalytic or the oligomerization core residues largely abolished the inhibiting effect of nsp15 on exogenous protein expression [chMDA5(N), chMAVS, chIRF7, EGFP] (**Fig 2A**), suggesting that the EndoU activity and oligomerization of nsp15 are indeed indispensable for its inhibition on protein synthesis. In addition, the expression level of wild type nsp15 was lower than the levels of mutated nsp15 (**Fig 2A**), suggesting that the effect on cellular protein synthesis also affected the expression of nsp15 itself. These data confirm that the inhibitory effect of nsp15 on the cellular protein synthesis requires its EndoU activity and oligomeric structure. The expression levels of the oligomerization-deficient nsp15 (D285A, D315A) were lower than those of the catalytic-deficient nsp15, suggesting that the inability to oligomerize may affect protein stability. Although far less profound than observed for wild type nsp15, the H238A mutation also resulted in a somewhat lower expression of co-transfected chIRF7, and the H238A, D285A and D315A mutations resulted in a lower expression of co-transfected EGFP (**Fig 2A**), suggesting that inactivation of single catalytic/oligomerization domain might not be sufficient to completely abolish the effect of nsp15 on protein expression.

**Fig 2.**
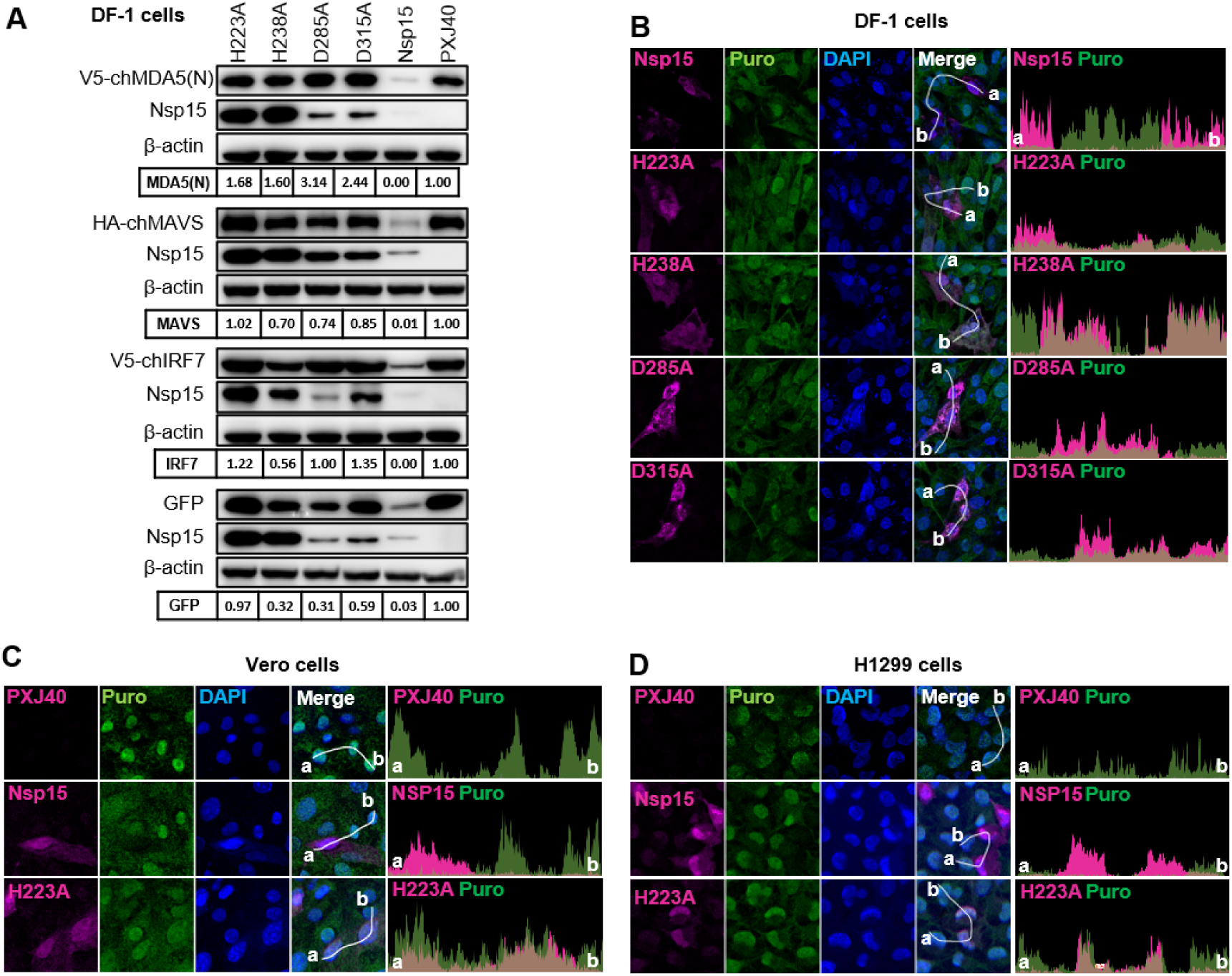
The catalytic activity and oligomeric structure of IBV nsp15 are indispensable for the inhibition of *de novo* protein synthesis in chicken cells as well as mammalian cells. (A) Plasmid encoding Flag-tagged catalytic-deficient nsp15 H223A, H238A, oligomerization-deficient nsp15 D285A, D315A, wild type nsp15, and the vector PXJ40, were each co-transfected with plasmids encoding V5-chMDA5(N), HA-chMAVS, V5-IRF7, or EGFP into DF-1 cells. After 24 h, Western blot analysis was performed using corresponding antibodies. β-actin was detected as loading control. Density of the bands was analysed by Image J, normalized to the signal of β-actin, and the ratio was presented relative to the density detected in the corresponding PXJ40 transfected cells. (B) DF-1 (C) Vero and H1299 cells, were transfected with the plasmid encoding wild type or mutated nsp15 and treated with puromycin (5 µg/ml) for 1 h at 23 h post-transfection (h.p.t), to label the *de novo* synthesized peptides. Indirect immunofluorescence was performed to detect nsp15 (magenta), puromycin (green), and nuclei (DAPI, blue). Fluorescence intensity of nsp15 and puromycin signal along the white line (from a to b) is indicated in the right panel (histogram plot).

We then examined the effect of wild type and mutated nsp15 on endogenous *de novo* protein synthesis by analysing the fluorescence intensity of the puromycin-labelled peptides signals. Indirect immunofluorescence analysis in DF-1 cells showed that, compared to the cells not expressing nsp15, wild type nsp15 expressing cells displayed a strongly reduced puromycin signal ; however, this was not observed in cells expressing mutated nsp15 (**Fig 2B**), demonstrating that EndoU activity and oligomerization structure are indeed indispensable for the inhibiting effect on cellular protein synthesis. We further assessed the inhibiting effect of nsp15 on cellular protein synthesis in Vero cells and H1299 cells, which are permissive cell lines for the IBV-Beaudette strain. Consistent with the results in DF-1 cells, indirect immunofluorescence analysis showed that the presence of wild type nsp15, but not of catalytic-deficient H223A nsp15, led to lower puromycin labelling signal in Vero and H1299 cells (**Fig 2C**). Altogether, these results demonstrate that the inhibiting effect on exogenous and endogenous protein synthesis by IBV nsp15 is a generalized inhibitory effect on cellular protein synthesis, and not restricted to cell types.

### Inhibition of *de novo* protein synthesis is a conserved feature of nsp15 from different genera of coronaviruses

We previously reported on the conserved activity of catalytic histidine residues of nsp15 on inhibition of SG formation, including nsp15 from IBV, PEDV, TGEV, PDCoV, SARS-CoV-1, MERS-CoVs, and SARS-CoV-2 [29]. This prompted us to investigate whether the observed inhibitory effect of IBV nsp15 on protein synthesis is conserved among nsp15 from different genera of coronaviruses and whether such function is dependent on the EndoU catalytic activity. For this purpose, the expression plasmids for wild type and catalytic-deficient nsp15 from the above-mentioned coronaviruses were co-transfected with a plasmid encoding EGFP or IBV N in Vero cells. Western blot analysis revealed that wild type nsp15 of PEDV, TGEV, PDCoV and SARS-CoV-1 reduced the expression EGFP and IBV N, while catalytic-deficient nsp15 did not (**Fig 3A-D**). Conversely, nsp15 of MERS-CoV and SARS-CoV-2 did not show a pronounced effect on EGFP or IBV N expression (**Fig 3E-F**). These data suggest that, consistently with the data for IBV nsp15 (**Fig 1**), also nsp15 from most of the tested coronaviruses can suppress exogenous protein expression.

**Fig 3.**
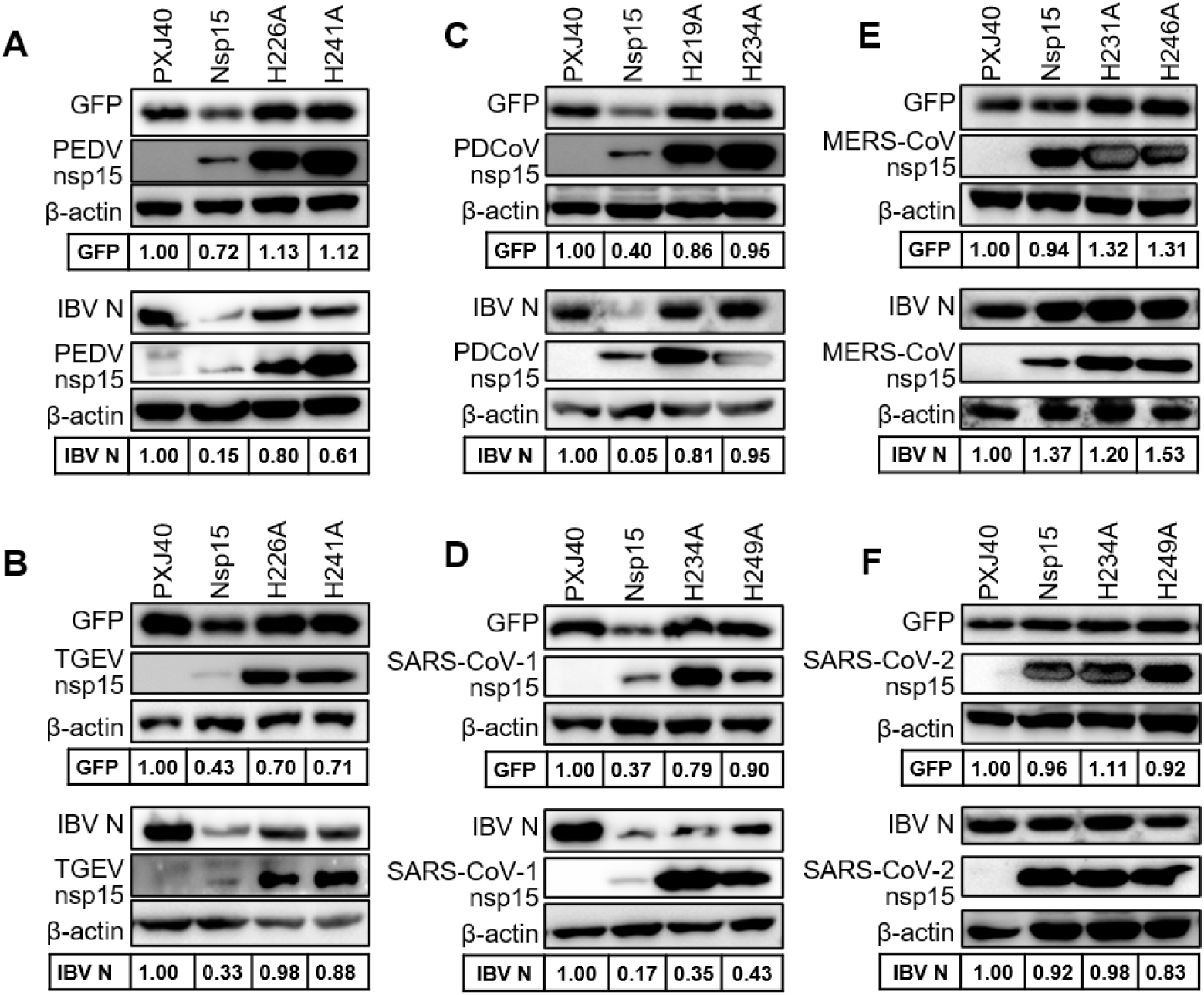
Nsp15 from most of coronaviruses inhibits expression of exogenous transfected plasmids. The plasmid encoding wild type or catalytic-deficient nsp15 from the indicated coronaviruses was co-transfected with the plasmid encoding EGFP or IBV N into Vero cells. After 24 h, Western blot analysis was performed using corresponding antibodies. β-actin was detected as loading control. Density of the bands of EGFP or IBV N were analysed by Image J, normalized to the signal of β-actin and presented relative to the PXJ40 group.

Next, we investigated whether nsp15 from different genera of coronaviruses also suppress endogenous protein expression. Consistent with the result on IBV nsp15 (**Fig 2B-D**), indirect immunofluorescence analysis revealed that PK15 cells expressing wild type nsp15 of PEDV, TGEV, PDCoV and HeLa cells expressing wild type nsp15 of SARS-CoV-1 displayed weaker puromycin labelling signals than cells not expressing nsp15 (**Fig 4**). This effect was largely abolished in catalytic-deficient nsp15 expressing cells (**Fig 4**). Consistent with the data in **Fig 3**, cells expressing nsp15 of MERS-CoV and nsp15 of SARS-CoV-2 also showed a reduction in puromycin signal compared to cells not expressing nsp15, but the effect was less pronounced than the reduction afforded by nsp15 from other coronaviruses, suggesting that nsp15 of MERS-CoV and SARS-CoV-2 might work somewhat differently from other coronaviruses nsp15. Similar effects of nsp15 on puromycin staining were observed when nsp15 of PEDV, TGEV, PDCoV were expressed in PK1, ST, and PK15 cells, respectively, and nsp15 of SARS-CoV-1, MERS-CoV and SARS-CoV-2 were expressed in Vero cells (**Supplementary Fig 2**), suggesting the inhibition of host protein translation by nsp15 is general and not restricted to specific cell types.

**Fig 4.**
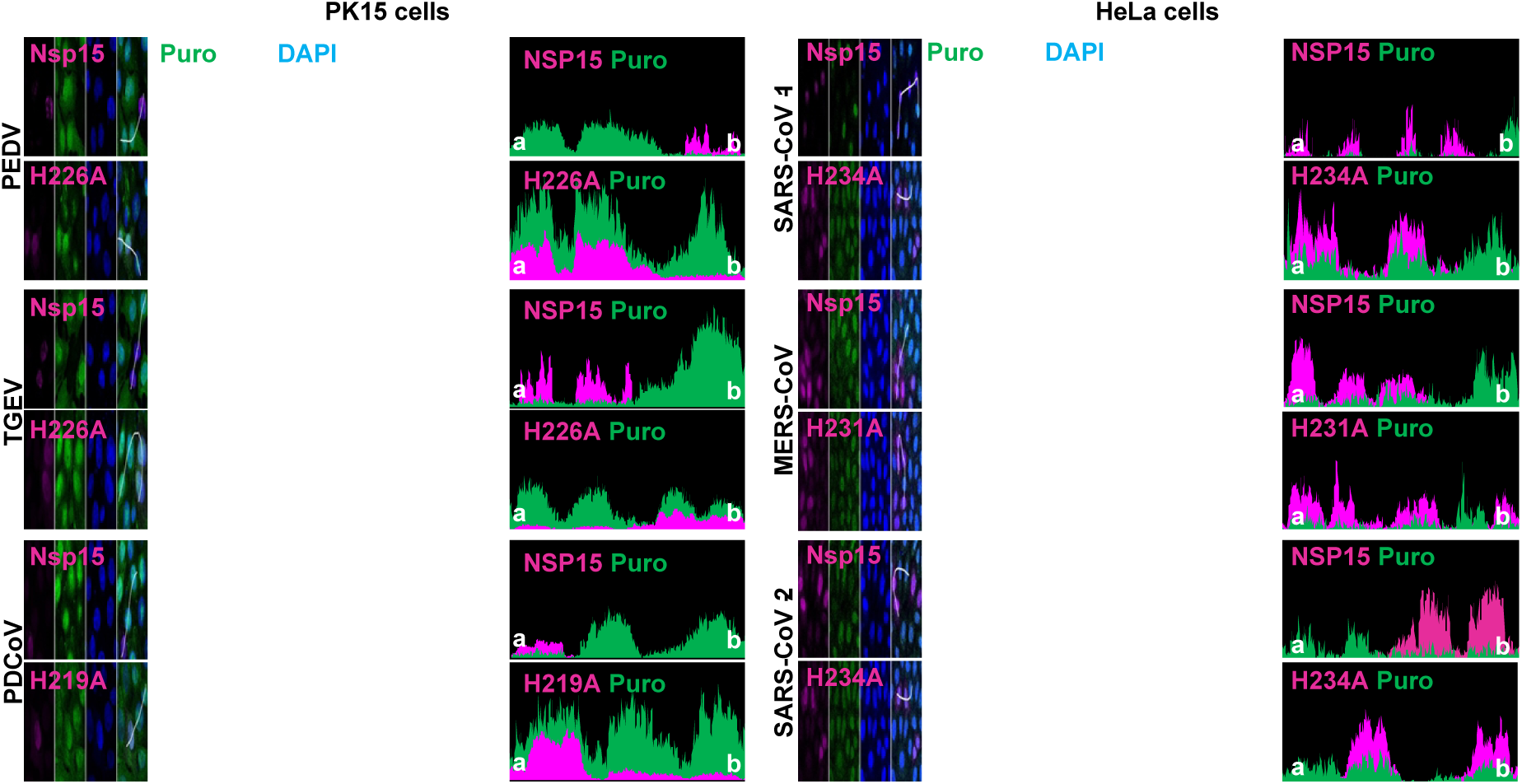
Nsp15 from different genera of coronavirus inhibits *de novo* protein synthesis. Porcine kidney 15 (PK15) cells were transfected with wild type nsp15 or the corresponding catalytic-deficient nsp15 from porcine coronaviruses PEDV, TGEV or PDCoV, human HeLa cells were transfected with wild type nsp15 or the corresponding catalytic-deficient nsp15 from human coronaviruses SARS-CoV-1, MERS-CoV or SARS-CoV-2. At 23 h.p.t, cells were treated with puromycin (5 µg/ml) for 1 h. Indirect immunofluorescence was performed using anti-Flag to detect nsp15 (magenta), anti-puromycin to detect puromycin-labelled *de novo* synthesized peptides (green), and DAPI to visualize nuclei (blue). Fluorescence intensity of nsp15 and puromycin signal along the white line (from a to b) is indicated in the right panel (histogram plot).

### Nsp15 from different genera of coronavirus alters the subcellular distribution of PABPC1 but does not significantly affect cellular mRNA localization

Considering that IBV nsp15 alone has a profound effect on *de novo* protein synthesis (**Fig 1 and Fig 2**) but that such effect is not mediated by direct degradation of mRNA (**Fig 1B**), we next investigated whether nsp15 may rather act on mRNA localization by for example preventing mRNA export from the nucleus to the cytoplasm leading to reduced synthesis of host proteins, as previously reported for Influenza virus NS1 protein [39]. mRNA shuttling is tightly regulated by RNA binding proteins such as PABPC1 [79–81]. We previously showed that under stress conditions [heat shock or sodium arsenite (ARS) treatment] the ability of nsp15 to relocate PABPC1 into the nucleus is a conserved feature among different genera of coronaviruses [29], we now ask whether this is accompanied by changes in mRNA distribution. We used a cross-reacting PABPC1 antibody that allowed us to visualize the localization of PABPC1 in DF-1 cells as well as mammalian cells, concomitantly to the distribution of mRNA by fluorescence *in situ* hybridization (FISH) using fluorescently labelled oligo (dT) probes. Consistent with our previous study, overexpression of IBV wild type nsp15, but not catalytic-deficient or oligomerization-deficient IBV nsp15, caused PABPC1 nuclear localization (yellow arrow) in some but not all nsp15-expressing DF-1 cells (**Fig 5A**), but this was not accompanied by accumulation of mRNA in the nucleus (red arrows), as otherwise observed in the positive control group treated with ARS (**Fig 5B**). The nuclear relocation of PABPC1 (yellow arrows) in nsp15-expressing cells was more obvious in Vero and H1299 cells than in DF-1 cells (**Fig 5B-D**), and again no obvious nuclear retention of mRNA (red arrows) was observed (**Fig 5B-D**). Closer observation revealed that in some cells, despite the lack of correlation between PABPC1 (yellow arrows) and mRNA distribution (red arrows), an apparent overlap between nsp15 or nsp15-H223A (white arrows) and mRNA (red arrows) could be observed in DF-1, H1299, and Vero cells (**Fig 5B**, **5D**, **supplementary Fig 3**), hinting at the possibility that nsp15 may bind to host mRNA.

**Fig 5.**
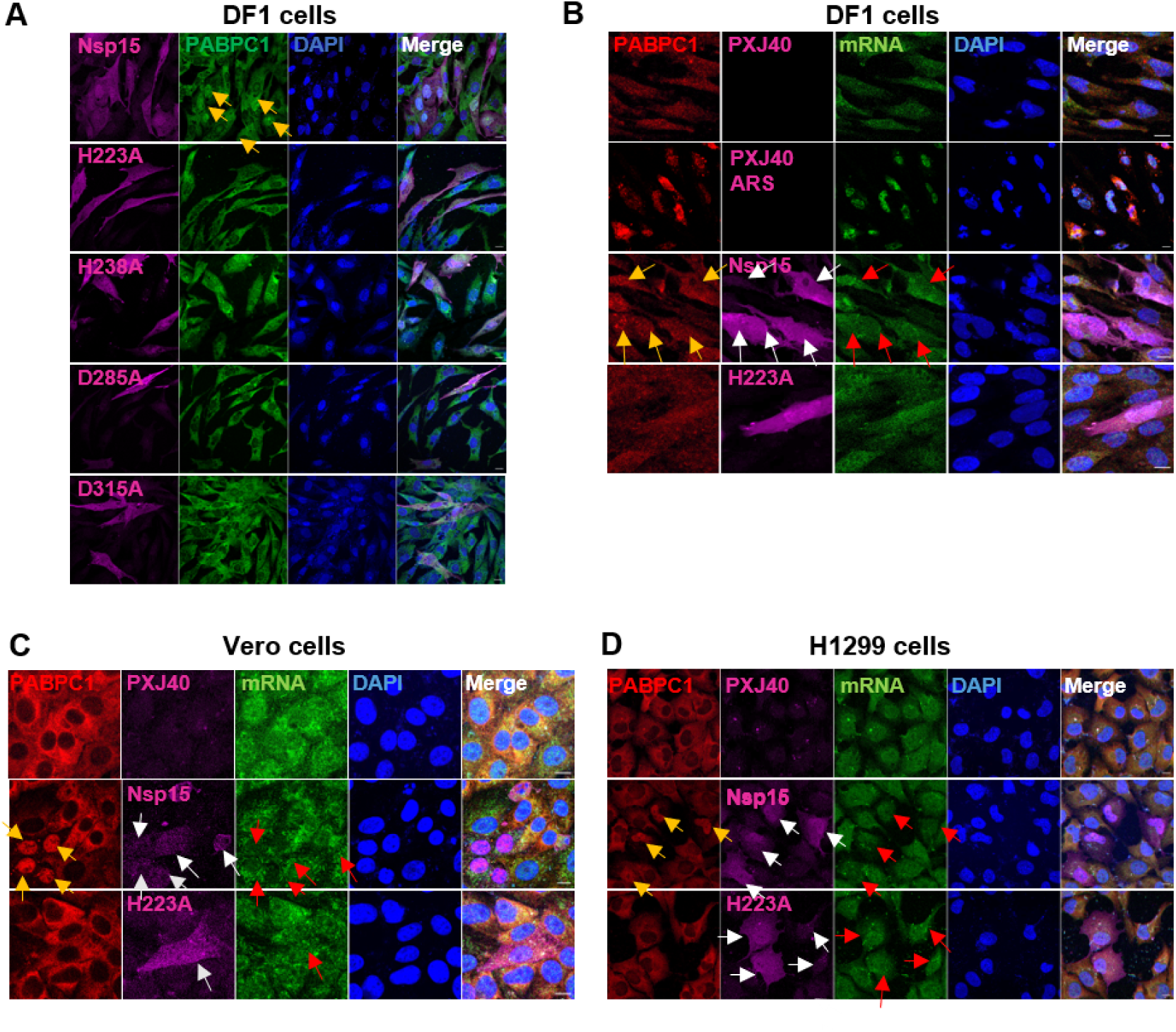
IBV nsp15 alters the localization of PABPC1 but not that of cellular mRNA. (A) Plasmids encoding wild type IBV nsp15 or the catalytic/oligomerization-deficient nsp15 (H223A, H238A, D285A, D315A) were transfected into DF-1 cells. After 24 h, indirect immunofluorescence was performed to visualize nsp15 (magenta), PABPC1 (green), and nucleus (blue). White arrows indicate cells that express wild type nsp15 and have nuclear accumulation of PABPC1. (B-C) DF-1, Vero, or H1299 cells were transfected with plasmids encoding IBV nsp15 or catalytic-deficient nsp15 H223A, or the empty PXJ40. After 24 h, *in situ* hybridization of mRNA was performed using oligo dT probes (green) followed by indirect immunofluorescence to detect IBV nsp15 (magenta) and PABPC1 (red). Nuclei were labelled by DAPI (blue). White arrows indicate the cells that express IBV nsp15, red arrows indicate the distribution of mRNA, yellow arrows indicate the cells with PABPC1 nuclear relocation. Treatment with 1 mM ARS for 30 min was used as positive control for stimulation of PABPC1 and mRNA nuclear localization.

After observing that the inhibitory effect on cellular protein synthesis is conserved among nsp15 from most coronaviruses (**Fig 3-4**), and that IBV nsp15 alters PABPC1 localization to nucleus without apparent effects on cellular mRNA distribution (**Fig 5**), we next investigated whether nuclear localization of PABPC1 in cells expressing nsp15 of PEDV, TGEV, PDCoV, SARS-CoV-1, MERS-CoV, or SARS-CoV-2 (as previously reported [29]) is accompanied by changes in mRNA distribution. In agreement with previous observation [29], also in PK15 cells and Vero cells, ARS treatment led to redistribution and colocalization of PABPC1 and mRNA to ARS induced cytoplasmic SGs (**Fig 6**). Whereas, similar to what was observed for IBV nsp15, overexpression of wild type, but not of the catalytic-deficient nsp15 of PEDV, TGEV and PDCoV, caused nuclear accumulation of PABPC1 (indicated by yellow arrows) but not of mRNA (indicated by red arrows) (**Fig 6A**). Interestingly, the mRNA signal was clearly lower (magenta arrows) in PEDV, TGEV and PDCoV nsp15 expressing cells that displayed nuclear relocation of PABPC1 (yellow arrows) than in cells that did not show this nuclear relocation, suggesting degradation of cellular mRNA. In Vero cells, wild type but not catalytic-deficient nsp15 of SARS-CoV-1led to nuclear localization of PABPC1 (yellow arrows) and this was not accompanied by a decrease or re-distribution of mRNA signal (red arrows) (**Fig 6B**). In line with the observed different effects of MERS-CoV and SARS-CoV-2 nsp15 on protein synthesis in **Fig 3**, **Fig 4, supplementary Fig 2**, only part of the cells expressing nsp15 of MERS-CoV and SARS-CoV-2 showed an altered localization of PABPC1 (yellow arrows) (**Fig 6B**); PABPC1 remained in cytoplasm in some nsp15 expressing cells (blue arrows). Again, in most nsp15-expressing cells, co-localization between nsp15 and mRNA signal or between nsp15-H238A and mRNA was observed (red arrows) (**Fig 6B**), suggesting binding of nsp15 to mRNA.

**Fig 6.**
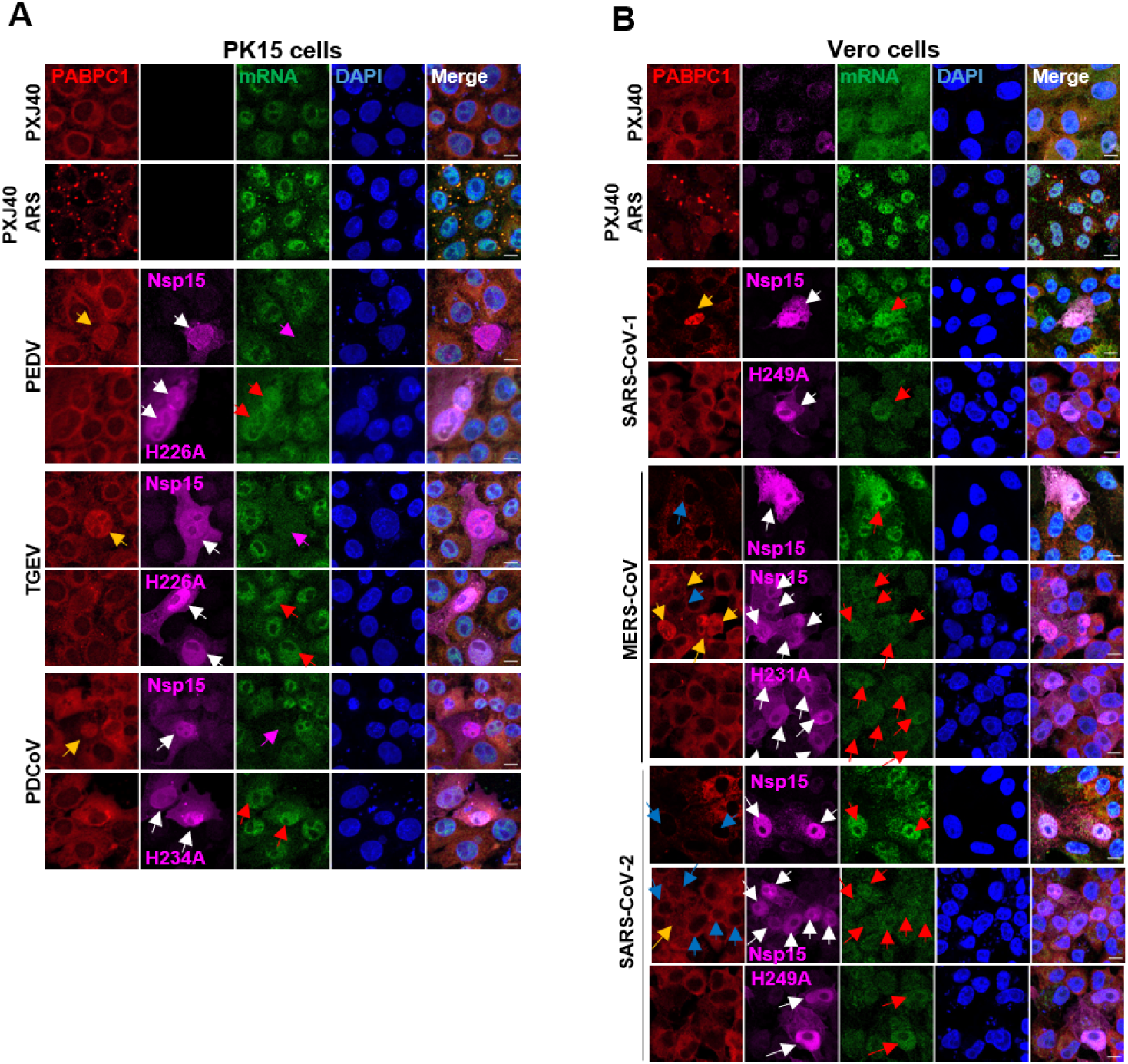
Nsp15 of PEDV, TGEV, PDCoV and SARS-CoV-1 alters the localization of PABPC1 but not that of cellular mRNA, nsp15 of MERS-CoV, SARS-CoV-2 does so in most but not all cells. Wild type nsp15 or catalytic-deficient nsp15 of the indicated coronaviruses was transfected into the indicated cell line. After 24 h, indirect immunofluorescence was performed to reveal the location of nsp15 (magenta), PABPC1 (green) and nucleus (blue). White arrows indicate the cells expressing nsp15, yellow arrows indicate the cells expressing nsp15 with a nuclear localization of PABPC1, blue arrows indicate the cells expressing nsp15 without nuclear localization of PABPC1, magenta arrows indicate the nsp15 expressing cells with weaker mRNA signal, red arrows indicate cells with overlapping of nsp15 and mRNA distribution. Treatment with 1 mM ARS for 30 min was used as positive control for stimulation of PABPC1 and mRNA nuclear localization.

Altogether these data suggest that subcellular redistribution to the nucleus of PABPC1 can be triggered by nsp15 of different genera of coronaviruses, and that this is dependent on the catalytic activity of nsp15. Furthermore, the PEDV, TGEV, or PDCoV nsp15-associated nuclear localisation of PABPC1 is accompanied by a weaker mRNA signal, possibly suggesting depletion or degradation of host mRNA by nsp15. Conversely, SARS-CoV-1, MERS-CoV and SARS-CoV-2 nsp15-mediated nuclear relocation of PABPC1 is not accompanied by changes in host mRNA distribution. Finally, overlap between nsp15 and mRNA signal was observed in some cells, suggesting nsp15 might competitively binds to host mRNA which in turns triggers relocation of PABPC1 into the nucleus.

### Nsp15 causes nuclear localization of PABPC1 accompanied by inhibition of cellular protein synthesis by targeting cytosolic factors

Next, we examined whether the nsp15-mediated PABPC1 nuclear relocation is associated with cellular protein synthesis shutoff. Overexpression of wild type, but not catalytic-deficient nsp15 from PEDV, TGEV, PDCoV, SARS-CoV-1 and IBV, altered the localization of PABPC1 (yellow arrows) and this was associated with a strong reduction in puromycin signal (**Fig 7**, red arrows). In agreement with the data in **Fig 6**, in cells expressing nsp15 of MERS-CoV and SARS-CoV-2, nuclear localization of PABPC1 was observed in some but not all cells (**Fig 7B**). In the MERS-CoV nsp15-expressing cells with nuclear PABPC1 localization, a weaker puromycin signal was observed (**Fig 7B**, red arrows), whereas in nsp15-expressing cells in which a change in localization of PABPC1 did not occur, no effect on puromycin signal was observed (**Fig 7B**, magenta arrows). In the SARS-CoV-2 nsp15 expressing cells, no obvious effect on puromycin signal was observed, no matter PABPC1 enters nucleus or retains in cytoplasm (**Fig 7**, magenta arrows). As expected, catalytic-deficient mutants of all nsp15 neither changed PABPC1 distribution, nor reduced the puromycin signal, demonstrating that the catalytic activity is required for both, PABPC1 redistribution and cellular translation shutoff. These results, combined with those in **Fig 2**, **Fig 4 and Fig 6**, demonstrate that overexpression of nsp15 triggers nuclear retention of PABPC1 and that this is associated with a strong inhibition of *de novo* protein synthesis. The SARS-CoV nsp15 is an exception for which does not obviously inhibits the *de novo* protein synthesis in Vero cells.

**Fig 7.**
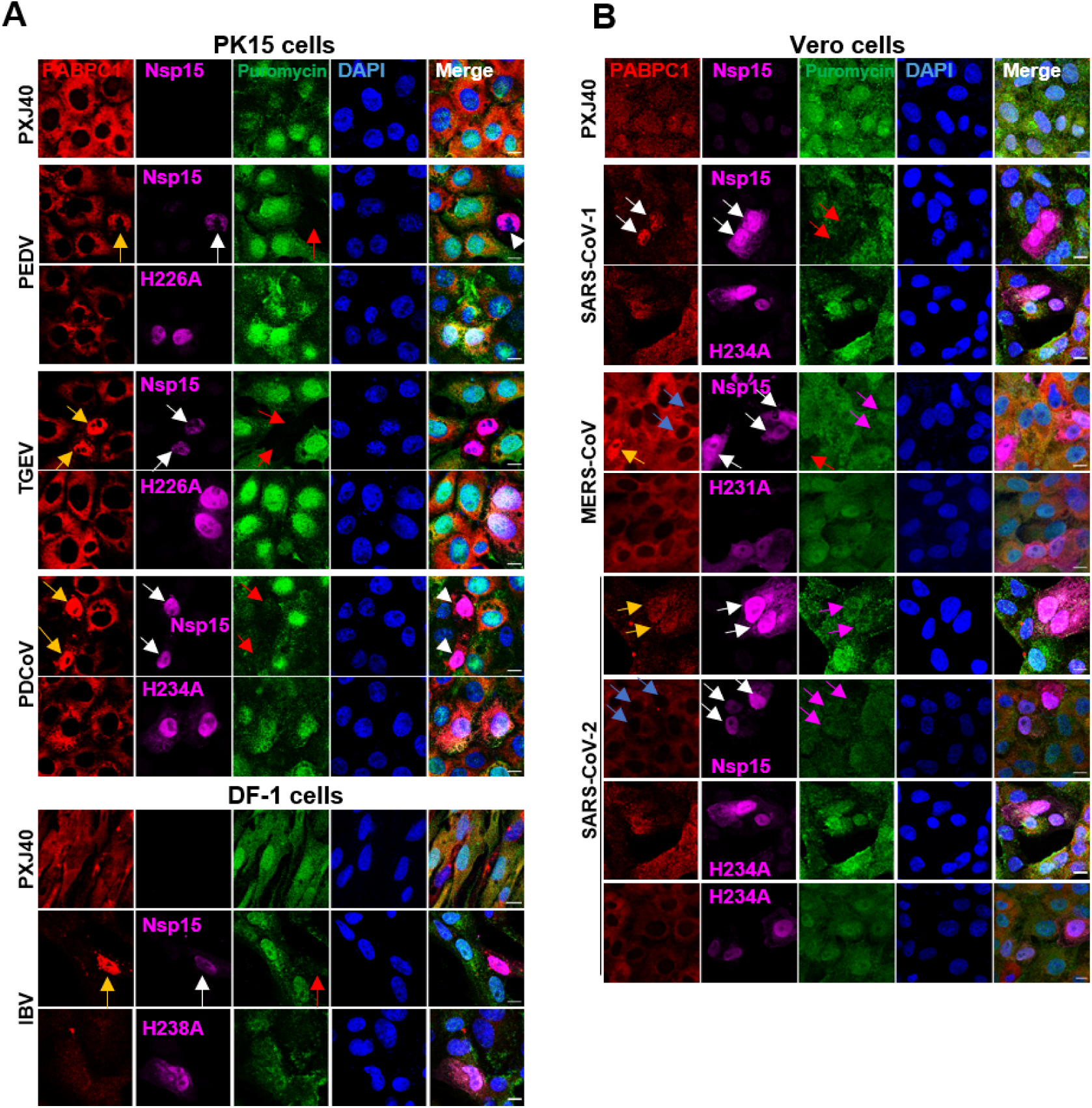
The nsp15-mediated relocation of PABPC1, is accompanied by inhibition of *de novo* protein synthesis. The indicated cell lines were transfected with PXJ40 or with a plasmid encoding wild type or the corresponding catalytic-deficient Flag-tagged nsp15 of the indicated coronaviruses. After 23 h, puromycin labelling (5 µg/ml) was performed for 1 h. Indirect immunofluorescence was performed using anti-Flag (magenta), anti-PABPC1 (red), and anti-puromycin (green) antibodies. The nuclei were stained with DAPI (blue). White arrows indicate nsp15 expressing cells, yellow arrows indicate nsp15-expressing cells with PABPC1 nuclear retention, blue arrows indicate nsp15-expressing cells without PABPC1 nuclear retention, red arrows indicating cells with PABPC1 nuclear retention and reduced puromycin labelling, magenta arrows indicate cells expressing nsp15 of MERS-CoV and SARS-CoV-2 in which puromycin labelling.

Next, we asked whether nsp15-induced protein translation shutoff may be caused by the ability of nsp15 to target cytosolic or nuclear factors or both. To this end, first a nucleus-free *in vitro* translation system (Rabbit Reticulocyte Lysate) was employed. Upon *in vitro* translation, IBV nsp15 reduced the expression of IBV N, IBV M, or luciferase, whereas catalytic-deficient nsp15 H223A and H238A did not (**Fig 8**). These data indicate that inhibition of translation by nsp15 can occur in the absence of nuclear factors, and that cytosolic factors involved in protein translation might be targeted by nsp15.

**Fig 8.**
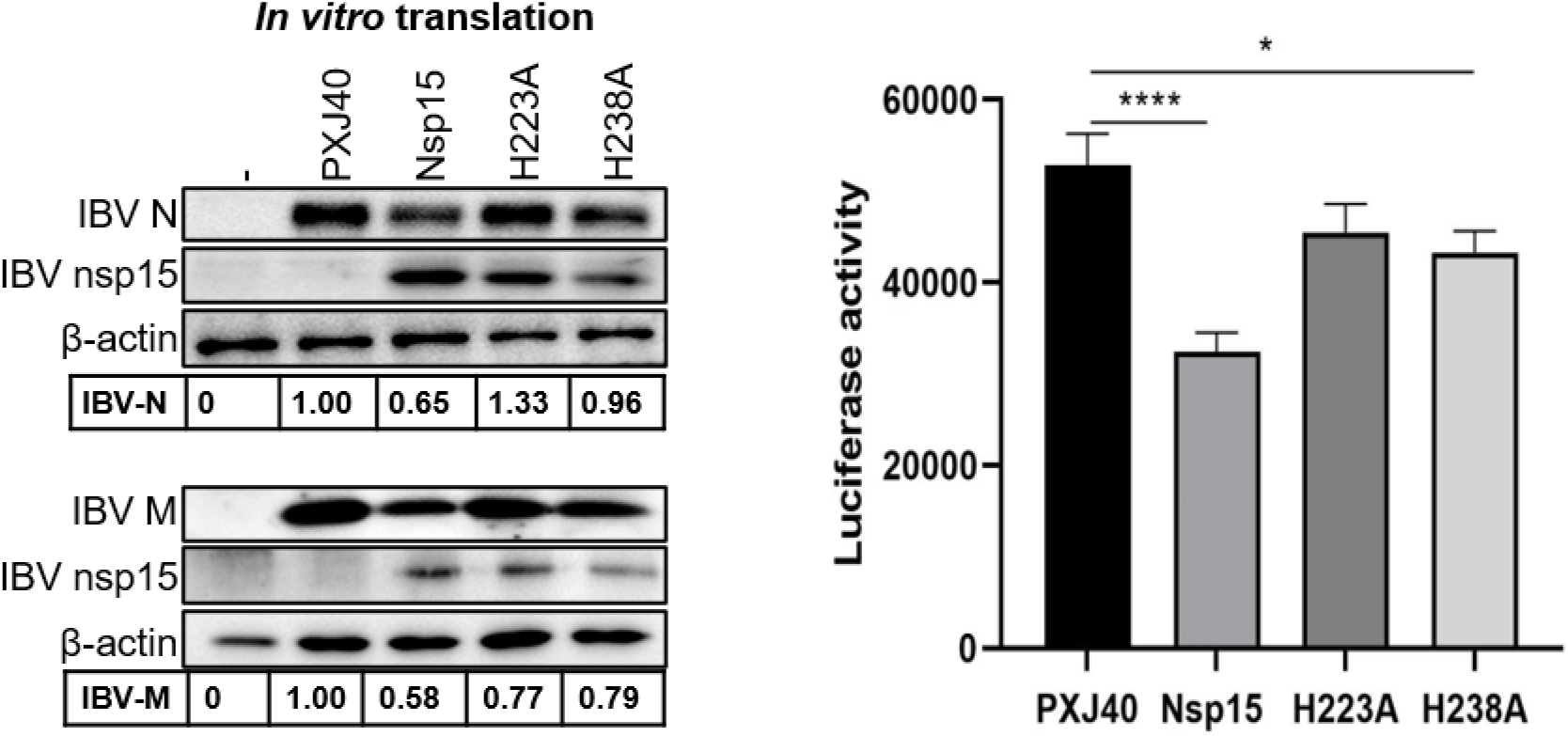
IBV nsp15 targets cytoplasmic as well as nuclear factors to inhibit protein translation. Plasmid encoding wild type or catalytic-deficient IBV nsp15 and reporter plasmid encoding IBV N or IBV M, or luciferase DNA, were co-incubated with Rabbit Reticulocyte Lysate for 1 h followed by Western blot analysis or luciferase assay. Density of the bands corresponding to the reporter proteins was normalized to the signal of β-actin and presented relative to the sample transfected with the empty vector PXJ40.

The observation that in nsp15-expressing cells PABPC1 nuclear retention is accompanied by protein shutoff but not by mRNA relocation, and that in some cells nsp15 localization largely overlaps with that of mRNA, combined with the observation that nsp15 may target cytosolic factors, suggests that nsp15 may interact with cellular mRNA itself or with (translation) complexes associated with the mRNA. The association of nsp15 to these complexes may compete or interfere with PABPC1 binding to the mRNA causing PABPC1 nuclear import, thereby leading to host protein translation shutoff. The targeting to host mRNA by nsp15 is further supported by the inability of catalytic-deficient nsp15 to trigger PABPC1 nuclear relocation and host-protein shutoff.

### Wild type IBV and the catalytic-deficient nsp15 mutant both induce host protein expression shutoff, but via different mechanisms

After having assessed that nsp15 may be involved in regulation of host protein translation, possibly through a mechanism involving targeting cytosolic mRNA and relocation of PABPC1, we next evaluated the effect of nsp15 on host protein translation in the context of a virus infection. To this end, we used the previously reported catalytic-deficient nsp15 recombinant IBV (rIBV-nsp15-H238A), with an Alanine substitution in the nsp15 catalytic domain H238 [29]. Western blot analysis after infection with wild type IBV (IBV-WT) or with the rIBV-nsp15-H238A mutant showed that, compared to uninfected cells, both viruses reduced puromycin labelling in DF-1 (**Fig 9A**) as well as H1299 cells (**Fig 9B**). IBV-WT did trigger a translational shutoff, however, it did not activate the PKR-eIF2α pathway, as dsRNA levels remained low until 24 h.p.i. and the phosphorylation levels of dsRNA sensor PKR and translation initiation factor eIF2α did not increase (**Fig 9B**), in agreement with our previous reports [29, 57, 82]. Thus, nsp15 helps virus degrade the viral dsRNA to escape the PKR-eIF2α dependent translation shutoff and subsequent stress response [29], which is detrimental for virus replication; in contrast, it might induce translation shut off through PKR-eIF2α independent strategies.

**Fig 9.**
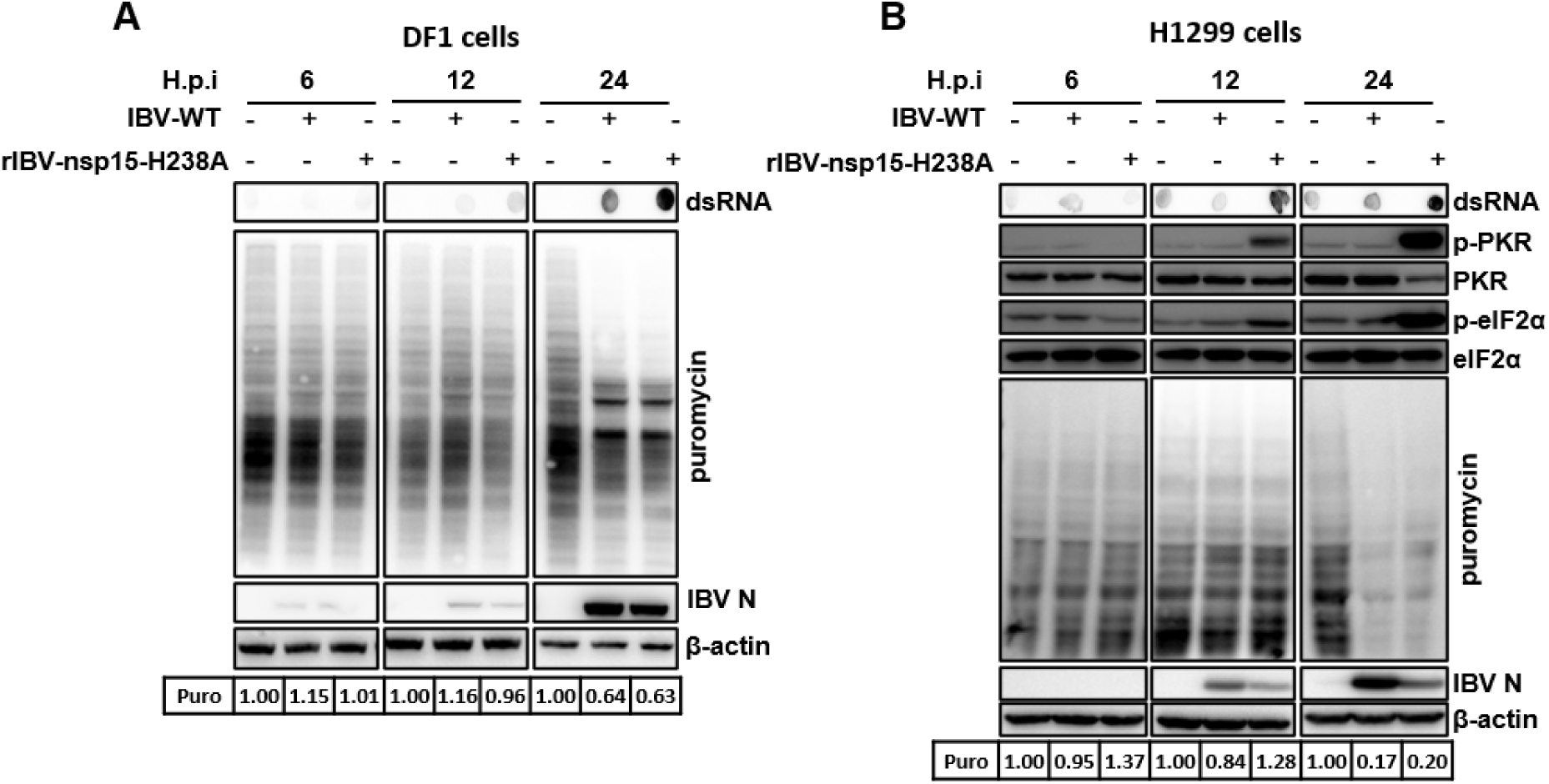
Both IBV-WT and rIBV-nsp15-H238A downregulate cellular protein synthesis but through different mechanisms. (A) DF-1 cells or (B) H1299 cells were infected with IBV-WT or rIBV-nsp15H1238A at an MOI of 1. At 6, 12, 24 h.p.i., cells were treated with puromycin (5 µg/ml) for 1 h, followed by western blot analysis to detect puromycin-labelled *de novo* peptides, IBV-N protein, and β-actin. Density of the puromycin labelled proteins was normalized to the signal of β-actin. Ratio of the puromycin–labelled *de novo* peptides of the infected cells (+) to that of the uninfected cells (-) at the same time h.p.i. is shown. (B) H1299 cells were infected as described above followed by dot blot analysis to detects dsRNA and western blot analysis to detect p-PKR, PKR, p-eIF2α, eIF2α.

Although rIBV-nsp15-H238A replicating less (assessed by the lower viral N protein expression), it reduced *de novo* protein synthesis to a similar extent as IBV-WT did (**Fig 9**). In agreement with previous report [29], infection with rIBV-nsp15-H238A leads to accumulation of higher levels of intracellular dsRNA intermediates than infection with IBV-WT, and this is accompanied by high levels of p-PKR as well as p-eIF2α (**Fig 9B**), ultimately leading to SGs formation and activation of the type I IFN response [29]. This, combined with our current data on puromycin labelling (**Fig 9**), confirms that rIBV-nsp15-H238A, but not IBV-WT, triggers a PKR-eIF2α-dependent translational shutoff that might impair both, host and viral protein translation initiation. This host-mediated translation initiation checkpoint shutoff triggered by rIBV-nsp15-H238A apparently did not benefit the virus, as synthesis of the viral N protein was lower compared to rIBV-WT. Therefore, the PKR-eIF2α-dependent translational shutoff, together with the induction of type I IFN [29], are responsible for the lower replication of rIBV-nsp15-H238A.

Taken all together, we hypothesize that both IBV-WT and rIBV-nsp15H238A cause host translation shutoff, but via different mechanisms: (1) IBV-WT controls the accumulation of viral dsRNA and therefore does not trigger the PKR-eIF2α pathway for >24h, but triggers a virus-mediate host translational shutoff that likely involves nsp15 and other viral proteins including E protein, the accessory protein 5a and 5b through a yet unidentified mechanism. (**Supplementary Fig 1, Fig 1**, **Fig 2**, **Fig 7**, **Fig 8**) [57]; (2) in the absence of nsp15 EndoU activity, rIBV-nsp15-H238A is unable to control the amount of viral dsRNA and induces translation shutoff through a host-mediated route involving the activation of the host PKR-eIF2α-mediated pathway [29].

### PABPC1 and mRNA nuclear relocation occurs upon rIBV-nsp15-H238A but not wild type IBV infection

As we observed after transfection a correlation between PABPC1 nuclear retention and inhibition of *de novo* protein synthesis only in cells overexpressing wild type but not catalytic-deficient nsp15 (**Fig 7**), while we observed a translation shutoff in both, IBV-WT- and rIBV-nsp15-H238A-infected cells (**Fig 9**), we next examined the localization of PABPC1 in cells infected with these two viruses. No changes in PABPC1 distribution were observed in most IBV-WT-infected cells at all time points after infection (**Fig 10A**, left panel), from 12 h.p.i. onwards, only few infected cells display the PABPC1 puncta aggregates colocalized with the SG core protein G3BP1 (**Supplementary Fig 4**), consistent with previous report [29]; in some but not all rIBV-nsp15-H238A-infected cells, from 12 h.p.i. onwards, nuclear localization of PABPC1 (**Fig 10A** right panel, **Supplementary Fig 5** right panel) or PABPC1 puncta aggregates colocalized with SG marker protein G3BP1 were observed (**Supplementary Fig 4, Supplementary Fig 5** right panel). As PABPC1 is involved in protein translation initiation in the cytoplasm, nuclear relocation or SG localization of PABPC1 after rIBV-nsp15-H238A infection, further suggests that rIBV-nsp15-H238A triggers a translation shutoff via a host-mediated mechanism. This host-mediated translation shutoff is detrimental to the virus as shown by the weaker viral N protein signal observed in rIBV-nsp15-H238A-infected cells with PABPC1 nuclear retention compared to infected cells without PABPC1 nuclear retention (**Fig 10A**, right panel, yellow cycles).

**Fig 10.**
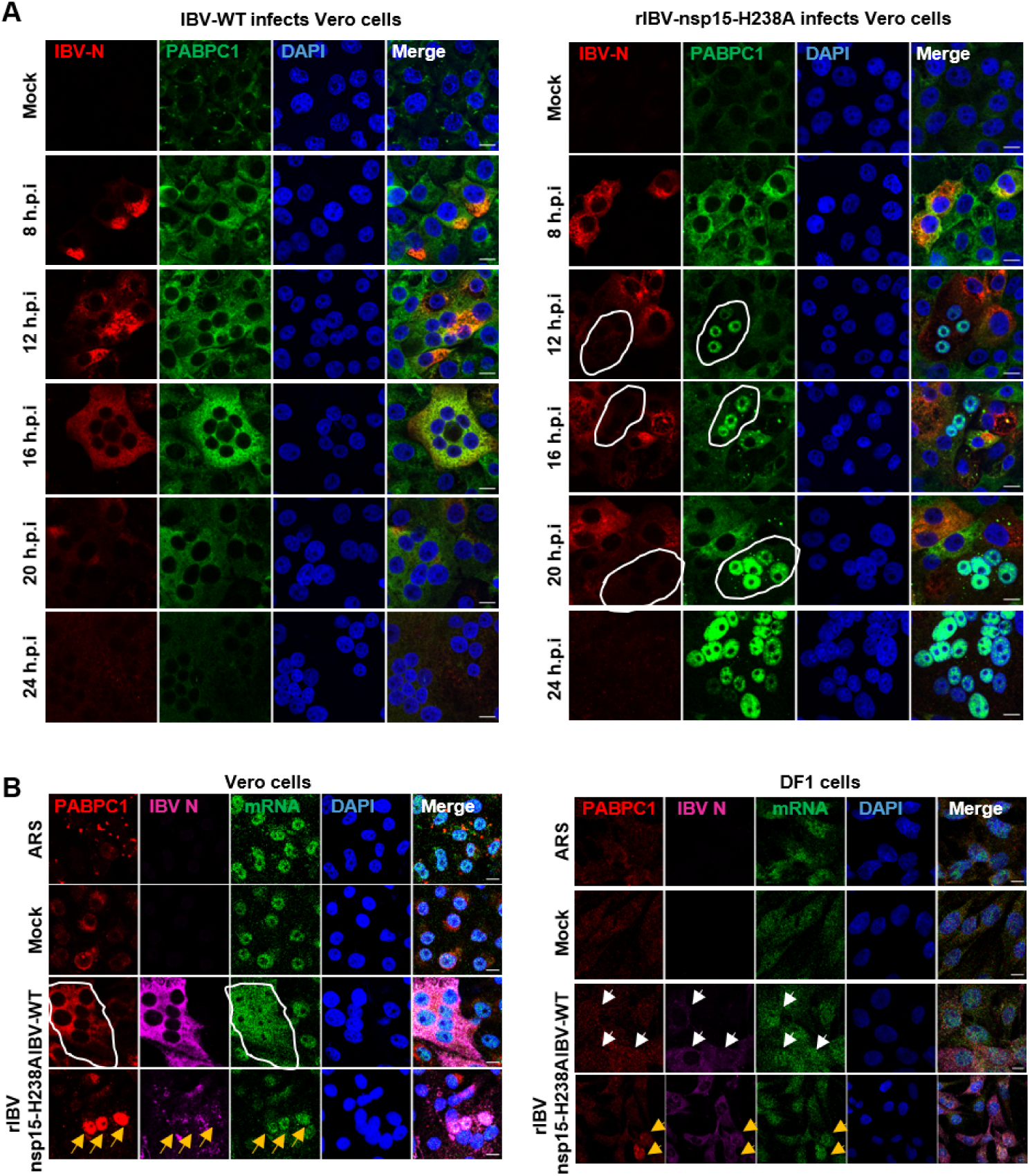
Infection with rIBV-nsp15-H238A, but not with IBV-WT, triggers nuclear retention of both, PABPC1 and mRNA and is associated with lower viral protein synthesis. (A) Vero cells were infected with IBV-WT (left panel) or rIBV-nsp15-H1238A (right panel) at an MOI of 1. At 8, 12, 16, 20 and 24 h.p.i, indirect immunofluorescence was performed to detect IBV-N protein (red), PABPC1 (green) and nuclei were stained with DAPI (blue). White circles indicate the rIBV-nsp15-H238A-infected cells that show nuclear localization of PABPC1 and weaker IBV-N signal compared to rIBV-nsp15-H238A-infected cells that do not display nuclear localization of PABPC1. (B) Vero and DF-1 cells were infected with IBV-WT or rIBV-nsp15-H1238A at an MOI of 1. At 16 h.p.i., FISH and indirect immunofluorescence were performed to detected PABPC1(red), IBV N (Purple), and mRNA (green). Nuclei were stained with DAPI (blue). White circle and white arrows indicate IBV-WT-infected cells in which no relocation of PABPC1 and a homogeneous distribution of mRNA was observed; yellow arrows indicate rIBV-nsp15-H238A-infected cells that show nuclear localization of both, PABPC1 and mRNA.

FISH analysis further showed that upon infection with IBV-WT in both, Vero and DF-1 cells, most mRNA is homogeneously distributed throughout the cells (**Fig 10B**, white circle or white arrows, **Supplementary Fig 4** left panel), whereas infection with rIBV-nsp15-H238A triggers PABPC1 nuclear retention or SG localization, and this is accompanied by mRNA nuclear accumulation and aggregates to SGs (**Fig 10B**, yellow arrows, **Supplementary Fig 4** right panel). These results suggest that in IBV-WT-infected cells, viral mRNA translation occurs; the presence of viral mRNA causes PABPC1 to bind and be retained in the cytoplasm; in rIBV-nsp15-H238A-infected cells however, the host-mediated translation shutoff halts both, host and viral mRNA translation, thereby releasing PABPC1 from cytosolic host and viral mRNA, redirecting it to the nucleus; the nuclear relocation of PABPC1 further retains the host mRNA in the nucleus.

### IBV nsp15 exhibits different subcellular localization after plasmid transfection and IBV infection

Subcellular localization is one of the key determinants for proteins access to their interacting partners, which provides important clues about proteins’ function. Thus, we examined the subcellular localization of IBV nsp15 expressed upon plasmid transfection and during IBV infection in DF-1 cells. Indirect immunofluorescence showed that exogenous Flag-tagged nsp15 localized to the cytoplasm, and likely also the nucleus (**Fig 11A**, upper panel; also visible in **Fig 2**, **Fig 5, and Fig 7**); however, upon IBV-WT or rIBV-nsp15-H238A infection, perinuclear aggregates of nsp15 or nsp15-H238A can be observed (**Fig 11A**, middle and low panels). These results show that nsp15 has different subcellular localizations in the context of plasmid transfection and virus infection. We hypothesize that during virus infection, nsp15 usually interacts with other viral nsps or viral RNA and locates to RTC to help virus genome replication and reduce viral dsRNA load [29], whereas when overexpressed, it cannot associate to RTC (owing to the absence of viral RNA and of other viral proteins) and is able to distribute throughout the cytoplasm and nucleus. The co-localization of nsp15 and RTC core protein RdRp nsp12 in **Fig 11B** further suggests that nsp15 indeed localizes to RTC during virus infection.

**Fig 11.**
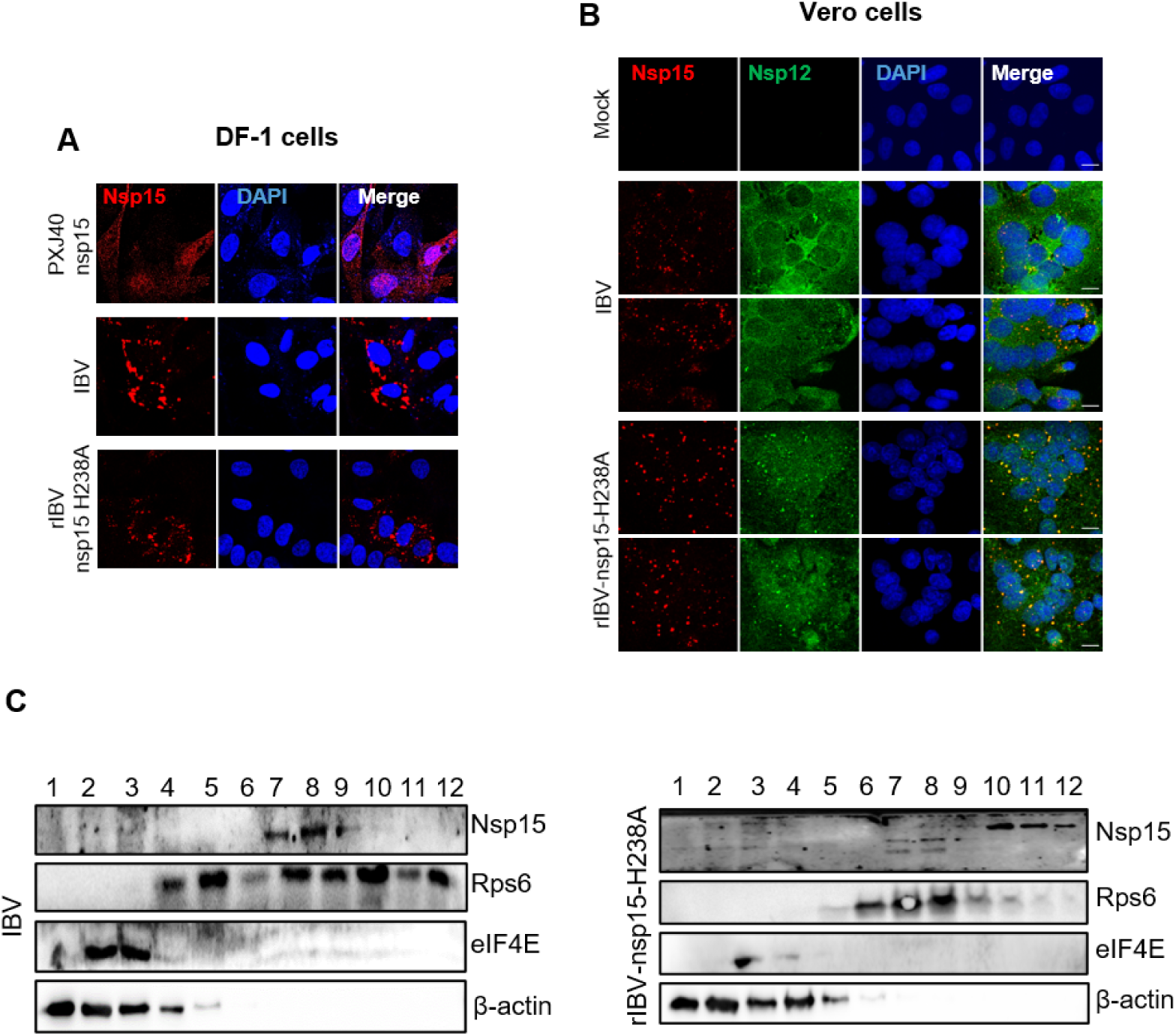
IBV nsp15 exhibits different subcellular localization after plasmid transfection and virus infection. (A) DF-1 cells were transfected with a plasmid coding Flag-tagged nsp15 (PXJ40F-nsp15). At 24 h.p.t, indirect immunofluorescence was performed with a chicken anti-Flag-tag antibody (red). Nuclei were stained with DAPI (blue). (B) Vero cells were infected with IBV-WT or rIBV-nsp15-H238A at an MOI=1. At 18 h.p.i, indirect immunofluorescence was performed with a mouse anti-IBV-nsp15 monoclonal antibody (red), a rabbit anti-IBV-nsp12 polyclonal antibody (green) and the nuclei were stained with DAPI (blue). (C) Vero cells were infected with IBV or rIBV-nsp15-H238A with an MOI of 1. At 18 h.p.i., cells were treated with 100 μg/mL cycloheximide (CHX) for 15 min at 37°C and subjected to 7–47% sucrose density gradient ultracentrifugation (38,000 rpm for 3 h), and the fractions were analysed by Western blot to detect nsp15, Rsp6, eIF4E, and β-actin (left panel).

We next asked whether nsp15 is associated to the translation machinery ribosomes and/or polysomes. For this purpose, Vero cells were infected with IBV-WT or rIBV-nsp15-H238A. After 18 h.p.i., cells were treated with translation elongation inhibitor cycloheximide (CHX) to immobilize ribosomes, lysed, and then subjected to a 7–47% sucrose density gradient ultracentrifugation. The fractions were analysed by Western blot to assess the distribution of nsp15 in the gradient fractions. The eIF4E protein was detected at fractions 1-4, and the ribosomal S6 protein (Rps6) was detected at fractions 4-11, representing mono-ribosomes and polysomes, respectively. In both, IBV-WT- and rIBV-nsp15-H238A-infected cells, nsp15 or nsp15-H238A was co-fractionated with Rsp6 at fraction 5-11, confirming its association with the host translation machinery, polysomes. It was noted that rIBV-nsp15-H238A infection led to a shift of Rsp6 to the lighter fractions 5-9, likely due to the host mediated translation shutoff (p-eIF2α-mediated translation initiation shutoff) which results in fewer polysomes and lower translation efficiency. It was worth noting that in rIBV-nsp15-H238A-infected cells, nsp15-H238A was detected in fractions 7-9, at the expected apparent molecular weight, whereas when detected in fractions 10-12, it displayed a higher apparent molecular weight, due to a yet unknown mechanism.

## Discussion

Several viruses encode ribonuclease and employ fine-tuned tactics to control viral as well as host mRNA expression balancing viral and host protein expression [73, 83–85]. The conserved EndoU nsp15 is the unique genetic marker of *Nidovirale*, as it is not present in other RNA virus families. In our previous study, nsp15 was found to suppress SG formation, either by preventing the accumulation of viral dsRNA or by targeting unknown host factors [29]. Thus, the role of nsp15 was shown to be more complex than originally believed, primarily because it has the potential to act on virus as well as cellular substrates. Although the role of nsp15 on the regulation of viral RNA is well studied [26, 27, 29, 86], the cellular substrates targeted by nsp15 are not yet known. In this study, by transfecting nsp15-encoding plasmids into eukaryotic cells, we show that nsp15 of different genera of coronaviruses inhibits the global protein synthesis, and that this inhibition is accompanied by the re-localization of the poly(A) tail binding protein PABPC1 to the nucleus. These activities were largely abolished when expressing catalytic-deficient nsp15, demonstrating the involvement of EndoU activity in the protein translation shutoff. This is the first report that coronavirus encoded EndoU is involved in regulation of host gene expression.

Protein translation initiation is a major step that determines the efficiency of host protein synthesis and viral protein synthesis [87]. PABPC1 is a nucleocytoplasmic shuttling protein that is predominantly located in the cytoplasm to help translation initiation by simultaneously interacting with the mRNA poly(A) tail and the eukaryotic initiation factor 4F complex (eIF4F), which binds to 5’ cap of mRNA and is part of the translation complex (TC); after bringing the mRNA-TC to ribosomes, PABPC1 is released and can bind the importin α/β complex, which mediates nuclear import of PABPC1 [80, 88–90]. In response to various pathogenic and non-pathogenic stressors, PABPC1 relocates to the nucleus or aggregates to the SGs [80, 91–93]. High levels of PABPC1 after nuclear relocation promote hyperadenylation and nuclear retention of mRNA [94], thereby restrict general gene expression.

In this study, we observed nuclear relocation of PABPC1 associated with protein translation shutoff in all nsp15 over-expressing cells. Considering that PABPC1 is able to shuttle between the nucleus and the cytoplasm [79] and that previous reports showed that blocking mRNA export from the nucleus to the cytoplasm usually causes nuclear retention of PABPC1 [80], we asked whether the nsp15 associated nuclear relocation of PABPC1 is the result of inhibition of PABPC1 nuclear export or enhanced nuclear import. *In situ* hybridization detecting poly(A) mRNA transcripts, did not reveal nuclear retention of mRNA in nsp15-expressing cells. This indicates that EndoU nsp15 does not block mRNA nuclear export to retain PABPC1 in the nucleus, and that nuclear relocation of PABPC1 is therefore the result of enhanced nuclear import. Nuclear import of PABPC1 is dependent on interaction with importin-α/β complex [80]. The motifs of PABPC1 that bind to importin-α/β complex are the same that recognize and bind mRNA [80, 95]; therefore, the dissociation from cytoplasmic mRNA leads to the exposure of PABPC1’s nuclear import signal and shuttling to the nucleus [80, 95], resulting in the observed inhibition of cellular protein expression. The dissociation of PABPC1 from cytoplasmic mRNA may be caused by the binding of nsp15 to (host) mRNA, thereby competing with PABPC1, as suggested by the general co-localization of nsp15 from all investigated coronaviruses with cellular mRNA. Furthermore, specifically in PK15 cells expressing nsp15 of porcine coronaviruses (TGEV, PEDV, PDCoV), a reduction in cellular mRNA signal was observed, supporting the possibility that nsp15 from some coronaviruses may not only bind, but also degrade host mRNA. Taken together, the relocation of PABPC1 to the nucleus, likely caused by the competition for cellular mRNA binding and/or mRNA degradation by nsp15, may therefore be the consequence, not the cause, of translation shutoff. This is further supported by the nucleus-free *in vitro* translation study, showing that inhibition of protein translation by nsp15 is not dependent on PABPC1 nucleus entry.

To assess whether nsp15 mediates host protein translation shutoff also during a virus infection, we infected cells with wild type IBV (IBV-WT) and the catalytic-deficient nsp15-H238A recombinant IBV (rIBV-nsp15-H238A). Interestingly, we found that both viruses inhibited host protein synthesis but via different mechanisms. IBV-WT inhibited host protein expression in an eIF2α check point-independent manner likely through a yet unknown virus-mediated mechanism, while rIBV-nsp15-H238A triggered host protein shutoff in a manner dependent on the activation of the dsRNA-PKR-eIF2α pathway, due to the accumulation of higher levels of dsRNA during this mutant virus infection. Surprisingly, nuclear retention of PABPC1 typical of nsp15-expressing cells, was not observed in IBV-WT-infected cells, whereas PABPC1 nucleus accumulation or SG aggregation was clearly observed in rIBV-nsp15-H238A-infected cells. The apparent contradiction with respect to PABPC1 re-localization between transfection and infection conditions might be explained by the presence of viral RNA during infection. Infection with IBV-WT causes host protein translation shutoff, but viral mRNAs (with poly(A) tail) may still bind to cytoplasmic PABPC1 for their own translation [96], thereby retaining PABPC1 in the cytoplasm. Infection with rIBV-nsp15-H238A, however, results in the accumulation of dsRNA due to the loss of nsp15 EndoU activity; this causes the activation of the PKR-eIF2α pathway, which in turn causes both, global mRNAs translation shuts off and aggregation of host and/or viral mRNAs to SGs, after which PABPC1 is released from host/viral mRNA and shuttles to the nucleus. This hypothesis is supported by previous studies showing the low abundance of cytoplasmic mRNA releases PABPC1 into nucleus [62, 95, 97]. Intriguingly, in rIBV-nsp15-H238A-infected cells, PABPC1 nuclear localization was accompanied by lower IBV-N protein signal compared to cells without PABPC1 nuclear relocation. This might be due to the stalled viral mRNA translation that releases more PABPC1 which in turn can enter into the nucleus. This phenomenon further suggests that re-localization of PABPC1 to the nucleus is a consequence of stalled translation initiation and dissociation of PABPC1 from mRNA.

The eIF2α phosphorylation-induced host shutoff inhibits both host and viral protein synthesis and is an anti-viral defence mechanism of the host [98, 99], while the eIF2α-independent translation shutoff might specifically regulate the host protein expression [70]. Thus, wild type IBV might specifically shutoff host protein expression by hijacking the host translation machinery through a yet unknown mechanism, in which nsp15, 5a, 5b, E, and S might be involved in (supplementary Fig 1) [56, 57]. Kint et al showed that the 5b is indispensable for host translation shut off by using the 5b null IBV [57], and Xiao demonstrated that IBV and SARS-CoV S protein bind to eIF3F and this interaction led to the inhibition of translation of a reporter gene [56]. Thus, we speculate that nsp15 inhibits host translation together with other viral proteins. However, rIBV-nsp15-H238A may cause translation shutoff through a host-mediated mechanisms based on phosphorylation of eIF2α caused by accumulation of dsRNA. The high level of phospho-eIF2α, induced by dsRNA activation of the PKR-eIF2α pathway, hampers the translation initiation step (Met-tRNA recruiting). This effectively stops both, host and viral protein translation, resulting in lower viral replication reflected by the reduced expression of the viral N protein. Nevertheless, the various mechanisms of translation shutoff triggered with or without nsp15 EndoU activity during virus infection, indicate that nsp15 plays a role in regulating protein translation in a way that benefits viral replication: by promoting the host translation shutoff via targeting to cytoplasmic factors and by avoiding the activation eIF2α-dependent translation shutoff via reducing the dsRNA formation.

Subcellular localization is one of the determinants for proteins to properly exert their functions. When over-expressed, nsp15 has a dispersed cytoplasmic and nuclear localization as no viral RNA or viral proteins are present to interact with or to recruit it to RTC. The cytoplasmic and nuclear distribution, enable nsp15 to target cellular substrates in the cytoplasm and the nucleus. During virus infection however, IBV nsp15 (**Fig 11**) and MHV nsp15 [100] display a perinuclear aggregated localization, colocalized with RdRp nsp12, the catalytic centre of the RTC [101]. Endoplasmic reticulum (ER) membrane transformation termed DMV during SARS, MERS or MHV infection, or zippered ER and spherules single membranes during IBV infections [22-24, 102–104], and collectively referred to as membrane rearrangements, provide a viral RNA replication microenvironment by accommodating the RTC. The association of coronavirus nsp15 with RTC implies that this EndoU may locate inside ER membrane rearrangements, as also suggested by IBV nsp15 perinuclear localization during infection and its co-localization with RTC-associated proteins like nsp12. Inside membrane rearrangements, nsp15 EndoU may cleave viral RNAs to reduce dsRNA accumulation. The membrane rearrangements are connected to each other, there is contiguity with the membrane donor of the ER [24], and the ribosomes are associated with the outer membrane [105–107]. Recently, cryotomography revealed that DMVs of MHV contain membrane spanning structures, a hexameric, crown-shaped pore complex surrounding a central channel that would allow RNA and protein transport [108]. The nucleocapsid structure was visualized on the cytosolic side of the pore, suggesting that the RNA is encapsidated following the export from DMVs [108]. Thus, although nsp15 is associated with the RTC or membrane vesicles during virus infection, it is still possible for it to locate outside membrane vesicles and to target host mRNA or the ribosomes in the outer membrane of vesicles, thereby interfering with host translation and this was also confirmed by our fractionation studies showing that nsp15 is associated to the translation machinery, and specifically to polysomes. Investigation into the kinetics of nsp15 production and formation of membrane rearrangements is necessary to track the location of nsp15 and analyse its roles during virus infection. In addition, regulation by other viral proteins interacting with nsp15 can also be a way to fine-tune nsp15’s roles during virus infection. We attempted to investigated whether nsp15 specifically inhibits host mRNA translation or also suppresses viral mRNA translation, by constructing the plasmid containing viral genomic 5’ leader sequence, 5’ UTR (untranslated region), transcription regulation sequence, IBV N gene, to mimic viral subgenomic mRNA structure; however, the translation of this construct was still supressed by co-transfected with IBV nsp15, probably due to lacking the 3’ sequence downstream of N gene. We will further construct the plasmid containing 5’ end and 3’ end of viral subgemomic mRNA, to fully mimic viral mRNA structure, for checking whether nsp15 viral mRNA translation.

Inhibiting host antiviral gene expression is an important strategy for viruses to antagonize the host innate immune response. Previous studies have reported that coronaviruses shut down host translation by various mechanisms. *α-* and *β-*coronaviruses employ nsp1 to trigger host translation shutoff via multiple strategies [109]: interference with ribosomal function [44, 110, 111], endo-nucleolytic cleavage of 5′ capped non-viral mRNA that triggers its degradation [44, 48, 50, 68], interference with nucleocytoplasmic transport of host mRNA leading to its nuclear retention [45, 46], or halting translation of host mRNA by targeting mRNA derived from the nucleus [48]. SARS-CoV-2 nsp14 and nsp16 also inhibit host translation, through a mechanism in which nsp16 suppresses global mRNA splicing and prevents the production of mature host mRNA [112, 113]. We previously reported that IBV, a *γ*-coronavirus, which lacks nsp1, also inhibits host translation through a yet unknown mechanisms that likely involves accessory protein 5b [53]. In the current study, by comparing the activity of wild type and catalytic-deficient nsp15 in the context of a viral infection or when over-expressed alone in eukaryotic cells after transfection, we were able to show that, besides the well characterized activity of nsp15 on viral dsRNA, nsp15 of IBV and of other coronaviruses exerts additional functions by also targeting host substrates, ultimately leading to suppression of host protein synthesis. The role of nsp15 in the regulation of host and viral protein expression is summarized in the model shown in **Fig 12:** (0) under steady-state conditions, newly transcribed mRNA is bound by PABPC1 in the nucleus, the complex shuttles to the cytoplasm where it immediately interacts with elongation initiation factors (eIF) part of the translation complex (TC), which in turn recruit first the 40S and then the 60S ribosomal subunits to start translation; translation initiation releases the mRNA from PABPC1 which binds to importin-α/β complex to shuttle back into the nucleus, where the cycle begins again. (1) When expressed alone (after transfection of nsp15-coding plasmids), nsp15 targets host substrates involved in mRNA translation by possibly targeting mRNA itself and/or factors associated to the TC, leading to dissociation of PABPC1 from cellular mRNA, its re-localization to the nucleus; for all this, the EndoU activity of nsp15 is indispensable. (2) Wild type IBV induces a host translation shutoff that specifically restricts host and benefits viral protein synthesis by hijacking the translation machinery; for this, nsp15 and the previously reported accessory protein 5b, 5a and E in our screen data (supplementary Fig 1), might both be involved. (3) Catalytic-deficient-nsp15-IBV can no longer regulate the levels of viral dsRNA [29] and therefore triggers a dsRNA-PKR-eIF2α-mediated host shutoff that hampers both, host and viral mRNA translation, causes nuclear re-localization of PABPC1, and restricts viral replication. The perinuclear localization of nsp15 during IBV infection, suggest that it can be associated to membrane rearrangements (zipped ER or spherules) that would give it access not only to viral RNAs to regulate the abundance of dsRNA [22], but also ribosomes and/or host mRNA normally associated to the same membrane rearrangements.

**Fig 12.**
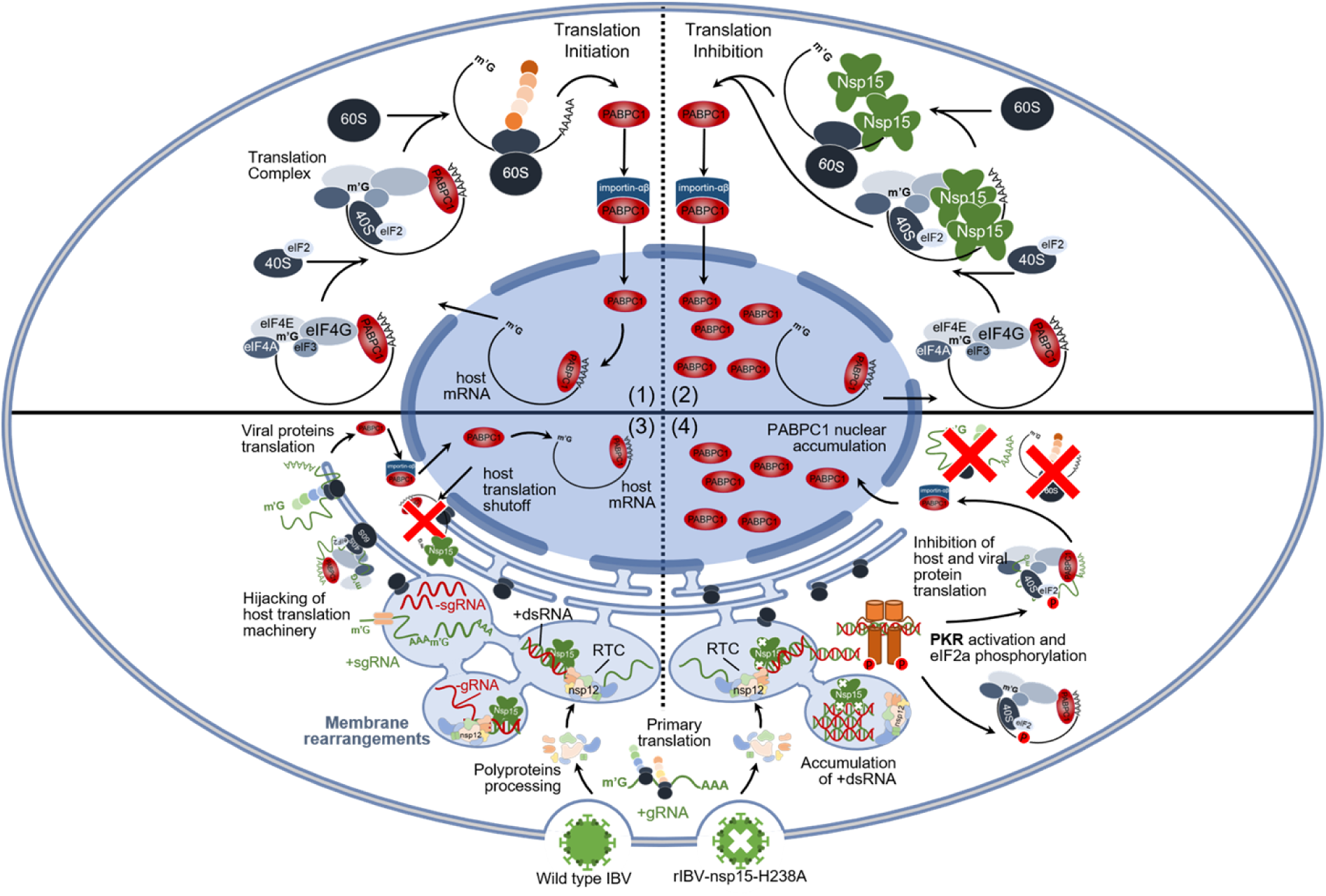
Working model of the mode of action on host protein translation of nsp15 when expressed alone or in the context of IBV infection. (1) host mRNA translation and PABPC1 turnover under normal conditions; (2) host mRNA translation upon overexpression of wild type IBV nsp15 leading to host-translation shutoff and nuclear accumulation of PABPC1; (3) simplified overview of the events occurring upon wild type IBV infection, ultimately leading to host-translation shutoff through a hijacking of the host translation machinery in favour of viral protein translation. (4) simplified overview of the events occurring upon catalytic mutant rIBV-nsp15-H238A infection. Catalytic-deficient nsp15 is no longer able to control the levels of viral dsRNA intermediates, leading to a dsRNA-PKR-eIF2α-mediated host-protein shutoff that limits both, host and viral protein synthesis, thereby leading to nuclear accumulation of PABPC1. For details refer to the main text.

This is a novel finding on the role of nsp15 to regulate host and viral gene expression, and increases our understanding on the regulation mechanisms of host translation by coronaviruses. Since the EndoU are conserved genetic marker of *Nidovirales,* the mechanisms of this EndoU as host translation suppressor may be a breakthrough in finding common strategies employed by coronaviruses, even by nidoviruses.

## Materials and Methods

### Cells and Viruses

Human non-small cell lung carcinoma H1299 cells were purchased from Cell Bank of Chinese Academy of Sciences (Shanghai, China). Chicken embryo fibroblasts DF-1 cells (ATCC® CRL-12203™), African green monkey kidney epithelial Vero cells (ATCC®CCL-81™) and human embryonic kidney HEK293T cells (ATCC® CRL-3216™) were purchased from ATCC. Porcine kidney epithelial cells (PK15) were provided from Prof. Hongjun Chen (Shanghai Veterinary Research Institute, CAAS, China). LLC-PK1 and ST cells were provided by Prof. Tongling Shan (Shanghai Academy of Agricultural Sciences, CAAS). H1299 cells were maintained in Roswell Park Memorial Institute 1640 medium (RPMI, 21875034, Gibco™) supplemented with 10% (v/v) foetal bovine serum (FBS, Gibco). The rest of the cell lines were grown in Dulbeco‘s modified eagle medium (DMEM, Gibco™) containing 10% fetal bovine serum (FBS, Gibco).

A mammalian cell adapted Beaudette IBV strain obtained from Prof Dingxiang Liu (Huanan Agricultural University, China) [114] was used in this study, as this IBV strain can be propagated in the DF-1 cells as well as in some mammalian cells, including Vero and H1299 cell lines [115]. The recombinant virus rIBV-nsp15-H238 was retrieved and its generation described in details our previous study [116].

### Plasmids

V5-tagged constitutively active form (N-terminus 1-1920 bp) of chicken MDA5 [V5-chMDA5(N)] and full-length chicken IRF7 (V5-chIRF7) were respectively cloned into pcDNA 3.1 vector and HA-tagged full-length chicken MAVS was cloned into the pCAGGS vector (provided by Yuqiang Cheng) [117]. The epitope tag is located at the C-terminus of the inserted gene. The plasmid pEGFP-N1 encoding enhanced green fluorescent protein (EGFP) was provided by Prof Yingjie Sun (Shanghai Veterinary Research Institute, CAAS, China). Construction of plasmids encoding IBV nsp2, nsp3, nsp4, nsp5, nsp6, nsp7, nsp8, nsp9, nsp10, nsp12, nsp13, nsp14, nsp15, nsp16, 3a, 3b, 5a, 5b, S, E, M, N, IAV NS1, PEDV nsp15, TGEV nsp15, SARS-CoV-1 nsp15, SARS-CoV-2 nsp15, and the catalytic-deficient mutants of the above nsp15 were cloned into vector PXJ40F, as described previously [29]. MERS-CoV nsp15 cDNA was purchased from Sangon Biotech, PDCoV-nsp15 cDNA was provided by Prof. Tongling Shan (Shanghai Veterinary Research Institute), and both were inserted into a PXJ40F vector. The oligomerization-deficient mutants of IBV nsp15 (IBV nsp15-D285A and IBV nsp15-D315A), and the catalytic-deficient mutants of MERS-CoV nsp15 and PDCoV nsp15 (MERS-CoV nsp15-H231A and PDCoV nsp15-H219A), were cloned using Mut Express II Fast Mutagenesis Kit V2 (C214, Vazyme). The mutagenesis primers were: for IBV nsp15-D285A, 5’-TGTTGT<u>ggc</u>TTTACTGCTTGATGATTTCTTAGAACTTC-3’ (F) and 5’-GCAGTAAA<u>gcc</u>ACAACAGTACACACTTGCTTGTAA-3’ (R); for IBV nsp15-D315A, 5’-GTGTCAATT<u>gct</u>TACCATAGCATAAATTTTATGACTTGG-3’ (F) and 5’-TGGTA<u>agc</u>AATTGACACTGTTACAACTTTTGACTT-3’ (R); for MERS-CoV nsp15-H231A, 5’-TTTTGAG<u>gcc</u>GTAGTCTATGGAGACTTCTCTCATACTACG-3’(F) and 5’-AGACTAC<u>ggc</u>CTCAAAAGCATAGTTTTCCAAGCC-3’(R); for PDCoV nsp15-H231A, 5’-CGGAACT<u>gcc</u>ACACTTATCTCACTAGTTAAAAACAAGTTTG-3’ (F) and 5’-TAAGTGT<u>ggc</u>AGTTCCGCCAATGACTGGACTG −3’ (R). The underlined sequences were the targeted sites for the mutations.

### Primary antibodies

Mouse anti-V5 (Thermo fisher scientific, #R961-25, horseradish peroxidase HRP-conjugated), mouse anti-HA (MBL, #M180-7, HRP-conjugated), mouse anti-Flag (MBL, #M185-7, HRP-conjugated), were diluted by 1:2500 for Western Blot; mouse anti-β-actin (CST, #3700S), rabbit anti-GFP (CST, #2956), chicken anti-Flag (Gentaur, #AFLAG), rabbit anti-phosphorylated PKR (Abcam, #ab32036), rabbit anti-PKR (CST, #12297), rabbit anti-phosphorylated eIF2α (CST, #3398), rabbit anti-eIF2α (CST, #5324), anti-RPS6 rabbit mAb (CST, #2217) and eIF4E rabbit mAb (CST, #2067), were diluted by 1:1000 dilution for Western Blot; rabbit anti-IBV N (provided by Prof Dingxiang Liu, South China Agricultural University, China) was diluted by 1:2000 for Western blot; rabbit anti-human PABPC1 (Abcam, #Ab21060, cross-reacts against chicken PABPC1 in DF-1 cells), rabbit anti-IBV-N, anti-IBV nsp12 rabbit pAb (provided by Prof Dingxiang Liu, South China Agricultural University, China), anti-IBV nsp15 mouse mAb (provided by Dr. Min Liao’s lab, Zhejiang University, China), were diluted by 1:500 for immunofluorescence; mouse anti-puromycin (Sigma-Aldrich, #MABE343) was diluted by 1:25000 for Western blot and 1:10000 for immunofluorescence; anti-dsRNA mouse mAb J2 (Scicons, #10010200) was diluted by 1:1000 for dot blot analysis.

### Secondary antibodies

Goat anti-rabbit IgG (H+L) (ABclonal, #AS014, HRP-conjugated) and goat anti-mouse IgG (H+L) (ABclonal, #AS003, HRP-conjugated) were diluted by 1:5000 for Western blot or dot blot; goat anti-chicken IgY (H+L) (Invitrogen, # A-11041, Alexa Fluor 568-conjugated), goat anti-mouse IgG (H+L) (Invitrogen, #A-11029, Alexa Fluor 488-conjugated), goat anti-rabbit IgG (H+L) (Invitrogen, #A-11034, Alexa Fluor 488-conjugated) were diluted by 1:500 for immunofluorescence.

### Chemicals

Puromycin (Merck, #58-58-2, reconstituted with sterile H_2_O to 50 mg/mL as stock solutions); Fugene (HD) (Promega, #E2311); Opti-MEM (Gibco™, #31985062); TRIzol regent (Life Technologies, #15596018); M-MLV (Promega, #M1701); Random primers (Invitrogen™, #48190011); SYBR green master mix (Dongsheng Biotech, #P2092); 4%-20% gradient SurePAGE gel (GenScript, #M00657); Tris-MOPS-SDS running buffer (GenScript, #M00138); Biotin-oligo d(T) (Promega, #Z5261, 0.2 μmol/L for mRNA FISH); Diethyl Pyrocarbonate (DEPC)-treated water (Invitrogen™, #4387937); Paraformaldehyde (Sigma-Aldrich, #158127); Triton X-100 (Sigma-Aldrich, #X100); BSA (Sigma-Aldrich, #A2153); SSC (Invitrogen™, #15557044); PBS (Sigma-Aldrich, #P3813); Dithiothreitol (DTT) (Thermo Scientific™, #R0861); RNase inhibitor (Promega, #N2611); Streptavidin (Invitrogen™, # SA1001, FITC - conjugated, 1:500 dilution for mRNA FISH); Diamidino-2-phenylindole (DAPI) (Thermo Scientific, #62247, 1:1000 dilution for IFA); Mounting medium (Sigma-Aldrich, #C9368); ARS (Sigma-Aldrich, #S7400).

### Plasmid transfection

Transient transfection using Fugene (HD) was described previously [29]. Fugene was used to transfect all cell lines employed in this study. Briefly, plasmid(s) and Fugene HD (M/V=1:3) were mixed in Opti-MEM. After incubation for 15 min at room temperature, the mixture was added to the cultured cells.

### Dual luciferase assay

Plasmids encoding IBV proteins (300 ng) (CMV promoter), human/chicken MDA5 (200 ng) (CMV promoter) or human/chicken MAVS (200 ng) (SV40 promoter), IFNβ promoter driven Firefly luciferase reporter (100 ng) and Renilla luciferase reporter pRL-TK (50 ng) (HSV TK promoter) were co-transfected into cells in a 24-well plate. To make the different IBV protein groups more comparable, the plasmids, except those encoding IBV proteins were mixed, and divided in equal volumes to which the plasmid coding each of the IBV protein was added in. Each co-transfection group was repeated twice. At 24 h.p.t., cells were lysed using the passive lysis buffer supplied by the Dual-Luciferase® Reporter Assay System (Promega, #E1910). Measurement of firefly luciferase activity by adding LAR II, and measurement of Renilla luciferase activity by adding Stop & Glo® were performed according to the manufacturer instructions using a luminometer reading (Cytation 5 imaging multimode reader, Biotek).

### Western blotting analysis

Briefly, cells transfected with plasmids for 24 h were lysed in lysis buffer and the cell lysates were resolved on 10% SDS-PAGE. To separate IBV-nsp7 (23 kDa), nsp8 (12 kDa), nsp9 (15 kDa), nsp12 (106 kDa), nsp15 (37 kDa) on the same gel, a commercial 4-20% acrylamide gradient SurePAGE gel was used. Different from SDS-PAGE gel electrophoresis, SurePAGE gel electrophoresis was performed using 1X Tris-MOPS-SDS running buffer. The proteins separated by the SDS-PAGE were transferred to a nitrocellulose membrane (GE life Sciences). The membrane was then incubated for 1 h at room temperature or overnight at 4 °C in blocking buffer (5% non-fat milk powder diluted in TBST (20 mM Tris, 150 mM NaCl, 0.1% Tween® 20 detergent), incubated with primary antibody diluted in blocking buffer, washed three times in TBST, incubated with HRP-conjugated secondary antibody diluted in blocking buffer, and again washed three times in TBST. Finally, signal was detected using a Tanon 4600 Chemiluminescent Imaging System (Bio Tanon, China) after development with luminol chemiluminescence reagent kit (Share-bio, China).

### Puromycin labelling

Puromycin resembles the 3′ end of tRNA and binds to growing peptide chains during translation, causing the stop of protein synthesis and release of premature polypeptides containing puromycin [78]. Cells were transfected with plasmid for 24 h or infected with IBV for indicated time, the cells were incubated with 5 µg/mL puromycin for 1 h at 37°C. The same amounts of 6 well plate cultured cells were lysed for western blotting analysis, or the cells cultured in 4 well chamber slide were fixed for indirect immunofluorescence assay.

### Sodium arsenite (ARS) treatment

Cells were seeded in six well culture plates and treated with 1 mM ARS for 30 min before being collected for mRNA *in situ* hybridization and indirect immunofluorescence analysis.

### Indirect immunofluorescence assay

Briefly, cells were fixed with 4% paraformaldehyde (diluted in PBS) for 15 min at room temperature (20-22°C), permeabilized with 0.5% Triton X-100 (diluted in PBS) for 15 min at room temperature, and incubated in blocking buffer (3% BSA diluted in PBS) for 1 h at 37°C. Cells were washed with PBS three times (5 min each) at the intervals of above steps on a shaker. Cells were then incubated with the primary antibody diluted in blocking buffer for 1 h at 37°C, followed by incubation with FITC- or TRITC-conjugated secondary antibody diluted in blocking buffer for 1 h at 37°C. In case of double staining, cells were then incubated with the other unconjugated primary antibody, followed by incubation with the corresponding FITC- or TRITC-conjugated secondary antibody. At the intervals of each incubation step, cells were washed three times (5 min each) with PBS buffer containing 0.2% Triton X-100 on a shaker. DAPI was then applied to stain the nuclei for 7 min at room temperature. Finally, cells were washed three times with PBS, mounted onto glass slides using prolong gold antifade mountant (Invitrogen), and examined by Zeiss LSM880 confocal microscope.

### Indirect immunofluorescence and mRNA fluorescence *in situ* hybridization (FISH)

To combine indirect immunofluorescence and mRNA FISH, cells were fixed for 15 min (4% paraformaldehyde in DEPC-treated PBS), permeabilized for 15 min (0.5% Triton X-100 in DEPC-treated PBS), and blocked for 1 h (3% BSA in DEPC-treated PBS), followed with endo-biotin blocking using a blocking kit (Invitrogen, #E21390) according to the manufacture instructions [118]. Cells were then incubated for 1 h at 37°C with the primary antibody. In case of double staining, the other primary antibody was then incubated in the same way. At the intervals of each incubation step, cells were washed three times with DEPC-treated PBS containing 0.2% Triton X-100. Cells were again fixed with 4% paraformaldehyde and washed three times with DEPC-treated PBS. Cells were then equilibrated in 2×SSC (1mg/mL t-RNA, 10% detran sulfate and 25% formamide) for 15 min at 42°C, followed by hybridization of biotin-oligo d(T) with the poly(A) tail of mRNA for approximately 12 h at 42°C in a humid environment. Biotin-oligo d(T) (0.2 μmol/L) were diluted in DEPC-treated PBS containing 0.2% TritonX-100, 1 mM DTT, and 200 units/ml RNase inhibitor. After the hybridization step, samples were washed with 2× SSC for 15 min and then with 0.5× SSC for 15 min, at 42°C on a shaker. Cells were again fixed with 4% paraformaldehyde and washed with DEPC-treated PBS. Cells were then incubated with Alexa Fluor-conjugated secondary antibodies for 30 min, and then with FITC-conjugated streptavidin for 30 min at 37°C. At the intervals of each step, cells were washed with DEPC-treated PBS containing 0.2% Triton X-100 three times. DAPI was then applied to stain the nuclei for 7 min at room temperature. Cells were washed again three times and mounted onto glass slides using mounting reagent. Cells were examined by Zeiss LSM880 confocal microscope.

### *In vitro* translation

0.5 μg of PXJ40-IBV-nsp15, 0.5 μg of reporter gene (PXJ40-IBV-N, PXJ40-IBV-M, or T7 luciferase control DNA), 40 μl TnT® Quick Master Mix (L1170, Promega), 1 μl Methionine (1 mM), nuclease-free water (to a final volume of 50 μl) were gently mixed by pipetting. The above mixture was then incubated for 90 min at 30°C. Samples of the translation reaction products were analysed by Western blot to detect protein expression level, or by luciferase assay to detect luciferase activity. For luciferase assay, 2.5 μl of translation reaction products and 50 μl of Luciferase Assay Reagent (Promega) were mixed by gently pipetting and subjected to luminometer reading (Cytation 5 imaging multimode reader, Biotek).

### Quantitative RT-PCR analysis

Total cellular RNAs were extracted using Trizol reagent. cDNAs were synthesized using M-MLV reverse transcriptase system. To prime the amplification of mRNAs in case their poly(A) tail is cleaved, random primers were used to synthesize cDNAs. The primers for quantitative PCR analysis were: for exogenous chicken *MDA5(N)*, 5’-AAAACGCAAGGAACGTGTCTG-3’ (F) and 5’-<u>GACCGAGGAGAGGGTTAGGG</u>-3’ (R); for exogenous chicken *MAVS*, 5’-ACATCCTTCCAGCTGTTGGC-3’ (F) and 5’-<u>CGTAATCTGGAACATCGTATGGG</u>-3’(R); for chicken *β-actin*, 5’-CCAGACATCAGGGTGTGATGG-3’ (F) and 5’-CTCCATATCATCCCAGTTGGTGA-3’(R). The underlined reverse primers target the V5 or HA epitope tag’s DNA sequence. In this way, mRNA transcribed from the plasmids were specifically amplified.

### dsRNA dot blot analysis

The accumulation of dsRNA was detected via dot blot analysis using an anti-dsRNA J2 antibody. Briefly, total cellular RNAs were extracted using Trizol reagent. RNA (2 μg) of each group was spotted on a Hybond-N+ membrane (GE Healthcare), followed by UV cross-linking (120 mJ/cm^2^) using SCIENTZ 03-II (Scientz Biotech). The membrane was then blocked in 5% non-fat milk dissolved in DEPC-treated TBS, incubated with mouse anti-dsRNA J2 antibody overnight at 4°C or for 1 h at room temperature, followed by incubation with the HRP conjugated goat anti-mouse secondary antibody for 1 h at room temperature. At the intervals of each incubation step, the membrane was washed three times with washing buffer (0.1% Tween® 20 detergent diluted in TBS). The dsRNA signals were detected using Tanon 4600 Chemiluminescent Imaging System after development with luminol chemiluminescence reagent kit.

### Polysome profile analysis

Vero cells were infected with 1 MOI of IBV or rIBV-nsp15-H238A for 16 h, then incubated with 100 μg/mL cycloheximide (CHX) (Selleck, Catalog No. S7418) for 15 min at 37°C to block translation elongation. Cells were washed and lysed by polysome extraction buffer (20 mM Tris-HCl pH 7.5, 100 mM KCl, 5 mM MgCl2, 0.5% Nonidet P-40) containing 100 μg/mL of CHX, protease inhibitors and RNase inhibitors. Lysates were clarified by centrifugation at 12,000×g for 15 min at 4°C, and supernatants were resolved on a linear sucrose gradient (7– 47% in buffer containing 20 mM Tris-Cl, pH 8.0, 140 mM KCl, 1.5 mM MgCl 2, 1 mM DTT, 1 mg/mL heparin) by centrifugation at 38,000 ×*g* at 4°C for 3 h. After centrifugation, each 1 ml fraction was collected and analysed by Western blot analysis.

### Densitometry

Image J program (NIH, USA) was used to quantify the intensities of corresponding bands of western blot, dsRNA dot blot, the intensity of puromycin signal in immunofluorescence image, and Pearson’s correlation coefficient of signals in immunofluorescence image.

### Statistical analysis

Data and statistical analysis were performed with Graphpad Prism 8 software. Significance was determined by ONE-Way ANOVA followed by Tukey’s post-hoc test. *p* < 0.05 was considered significant.

### Data availability statement

All relevant data are within the paper and its Supporting Information files.

## Acknowledgments

We would like to thank Prof. Dingxiang Liu (South China Agricultural University, China) for providing the IBV Beaudette strain and the rabbit anti-IBV-nsp12 polyclonal antibody, and for his excellent scientific advice. We also grateful to Prof. Bin Li (Jiangsu Academy of Agricultural Sciences, China) for providing TGEV nsp15 cDNA, Prof. Tongling Shan (Shanghai Academy of Agricultural Sciences, CAAS) for providing LLC-PK1 and ST cells, pDCoV cDNA, Dr. Min Liao (Zhejiang University, China) for providing IBV nsp15 monoclonal antibody.

## Disclosure

The authors have no financial conflict of interest.

## Grant support

This study was supported by the National Key Technologies Research and Development Program (No. 2021YFD1801104), the National Natural Science Foundation of China (32172834, 31772724).

## Author Contributions

**Conceptualization:** Ying Liao, Maria Forlenza, Xiaoqian Gong

**Formal analysis:** Ying Liao, Maria Forlenza, Xiaoqian Gong, Edwin Tijhaar, Shanhuan Feng

**Funding acquisition:** Ying Liao, Chan Ding

**Investigation:** Xiaoqian Gong, Shanhuan Feng, Bo Gao, Wenlian Weng, Hongyan Chu, Wenxiang Xue, Yanmei Yuan, Yuqiang Cheng

**Project Administration:** Ying Liao, Maria Forlenza

**Resource:** Cuiping Song, Lei Tan, Xusheng Qiu, Chan Ding, Min Liao

**Supervision:** Ying Liao, Maria Forlenza, Shouguo Fang, Chan Ding

**Writing-original draft:** Ying Liao, Maria Forlenza, Xiaoqian Gong, Edwin Tijhaar

**Writing-review & editing:** Ying Liao, Maria Forlenza, Edwin Tijhaar

---

**Supplementary Fig 1.**
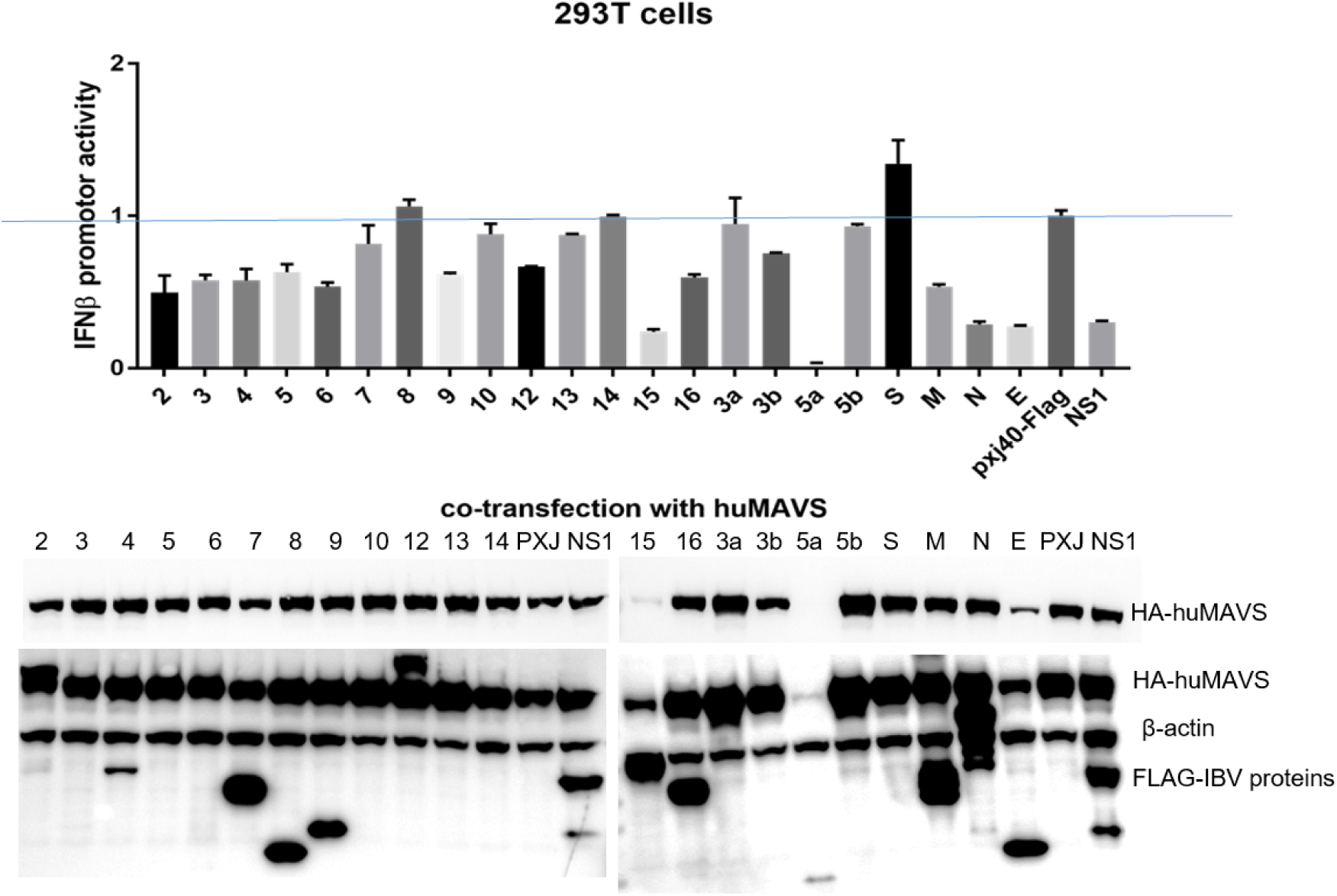
IBV nsp15 downregulates MAVS-mediated IFNβ induction and the expression of co-transfected plasmid. 293T cells were seeded in 48 well plates at 3×10^4^ cells/well (for luciferase assay) or 12 well plates at a density of 10^5^ cells/well (for Western blot). Cells were co-transfected with Flag-tagged IBV proteins (nsp2, nsp3, nsp4, nsp5, nsp6, nsp7, nsp8, nsp9, nsp10, nsp12, nsp13, nsp14, nsp15, nsp16, 3a, 3b, 5a, 5b, S, M, N, E) together with HA-huMAVS, the reporter plasmid encoding firefly luciferase driven by the inducible IFNβ promoter, and the control plasmid pRL-TK encoding Renilla luciferase driven by the constitutive HSV TK promotor. The PXJ40-Flag and IAV Flag-tagged NS1 were included in parallel experiment control. After 24 h, cells in 48 well plates were lysed, the firefly and Renilla luciferase activities were measured. The IFNβ promoter activity was normalized to Renilla and presented relative to the PXJ40-Flag control. Bars indicated the average of two co-transfection experiments performed independently. Cells in 12 well plates were lysed and subjected to Western blot to verify the protein expression. The membranes were first probed with an anti-HA antibody to detect huMAVS, followed by re-probing with an anti-Flag antibody to detect IBV proteins and IAV NS1, and then probed with an anti-actin antibody to detect actin as loading control.

**Supplementary Fig 2.**
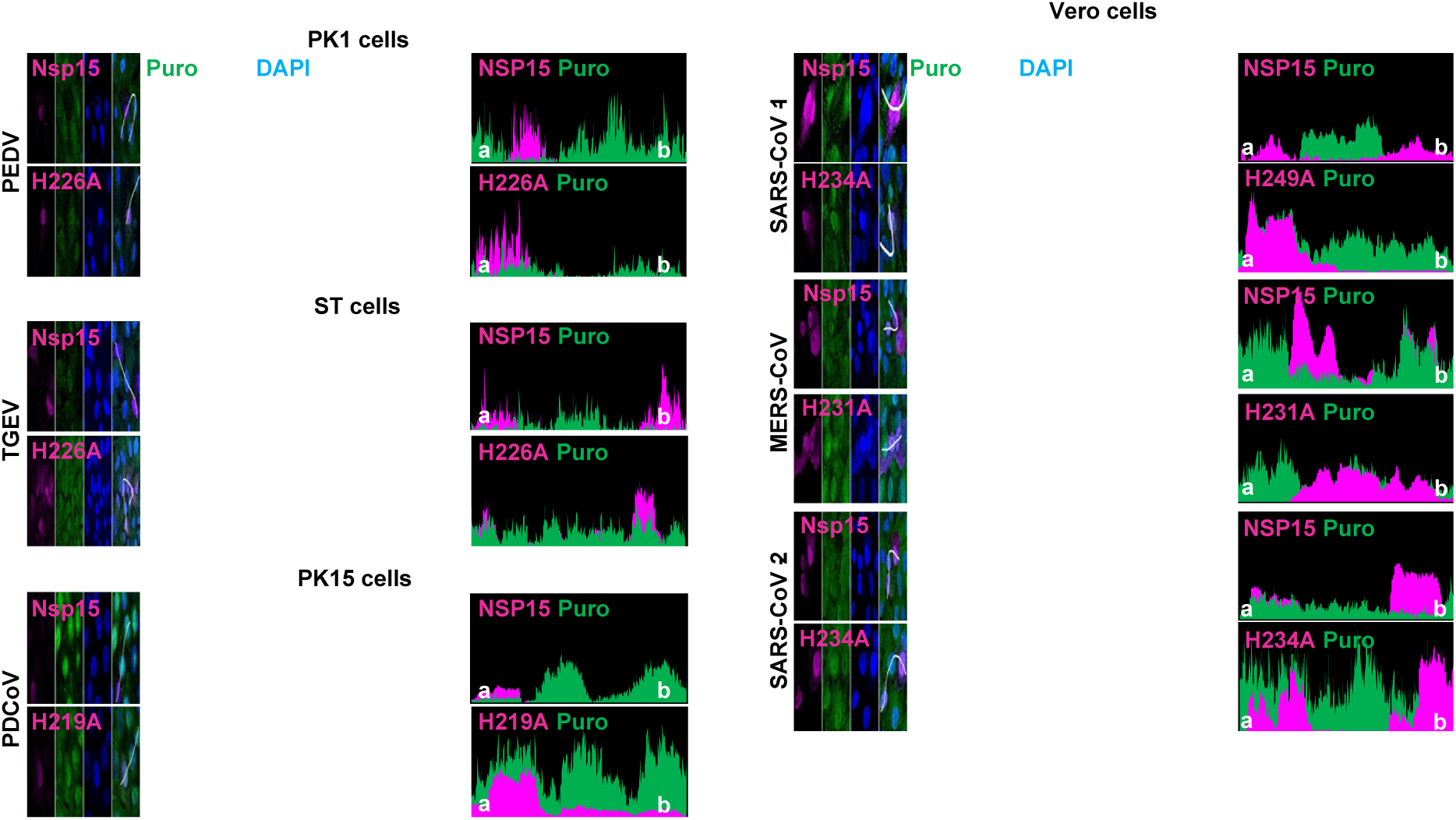
Nsp15 from different genera of coronavirus inhibits *de novo* protein synthesis. The porcine (PK1, ST and PK15) cells and Vero cells were transfected with wild type nsp15 or the corresponding catalytic-deficient nsp15 from the indicated coronaviruses. At 23 h.p.t, cells were treated with puromycin (5 µg/ml) for 1 h. Indirect immunofluorescence was performed using anti-Flag to detect nsp15 (magenta), anti-puromycin to detect puromycin-labelled *de novo* synthesized peptides (green), and DAPI to visualize nuclei (blue). Fluorescence intensity of nsp15 and puromycin signal along the white line (from a to b) is indicated in the right panel (histogram plot).

**Supplementary Fig 3.**
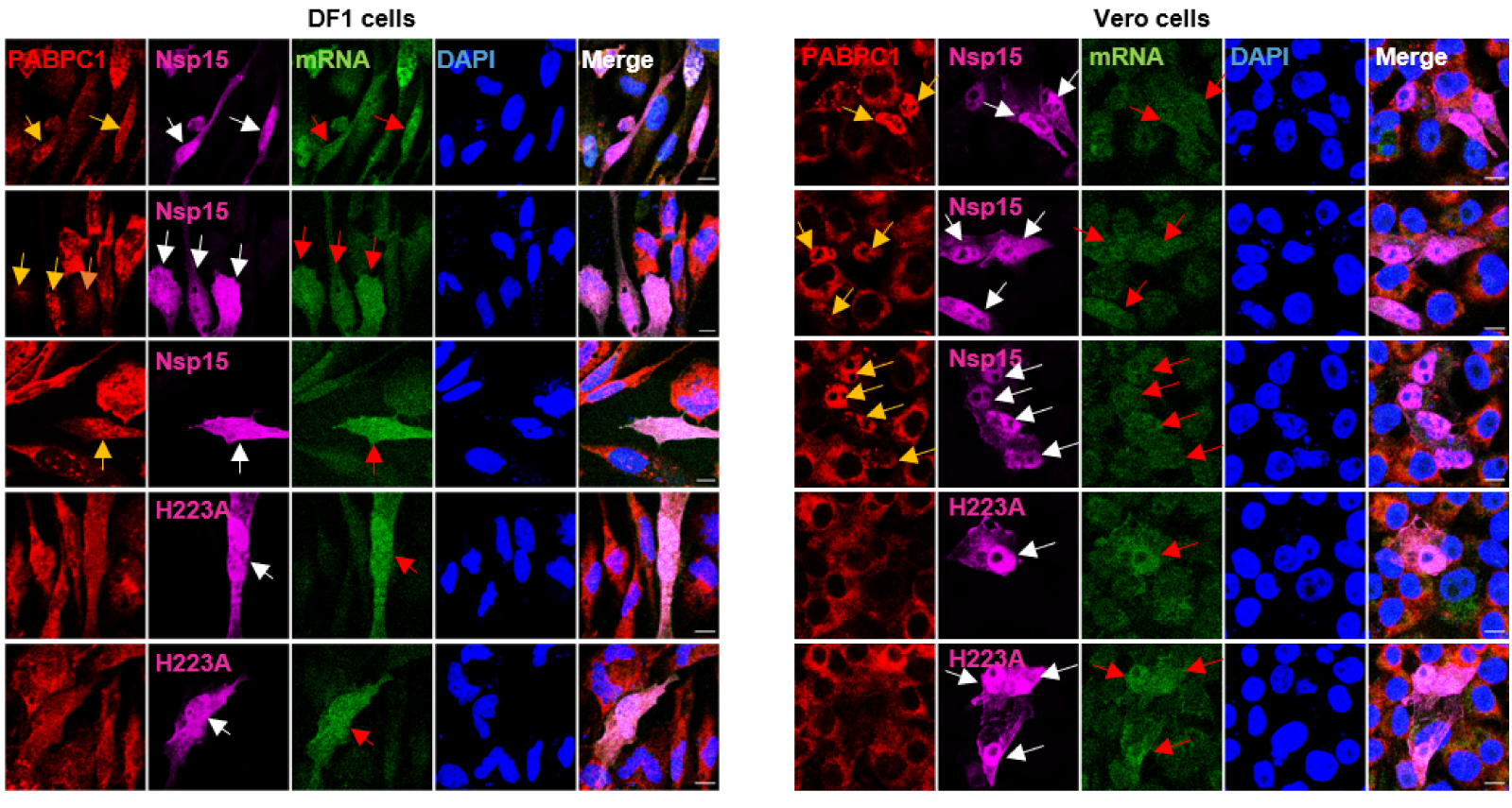
Distribution of IBV nsp15 overlaps with that of mRNA. Wild type IBV nsp15 or the catalytic/oligomerization-deficient nsp15 (H223A, H238A, D285A, D315A) was transfected into DF-1 or Vero cells. After 24 h, *in situ* hybridization followed by indirect immunofluorescence was performed to visualize PABPC1 (red), nsp15 (magenta), cellular mRNA (green) and nuclei (blue). Yellow arrows indicate cells that express wild type nsp15 with nuclear accumulation of PABPC1. White arrows and red arrows indicate cells expressing either wild type or mutated nsp15 in which location of nsp15 strongly overlaps with that of mRNA.

**Supplementary figure 4.**
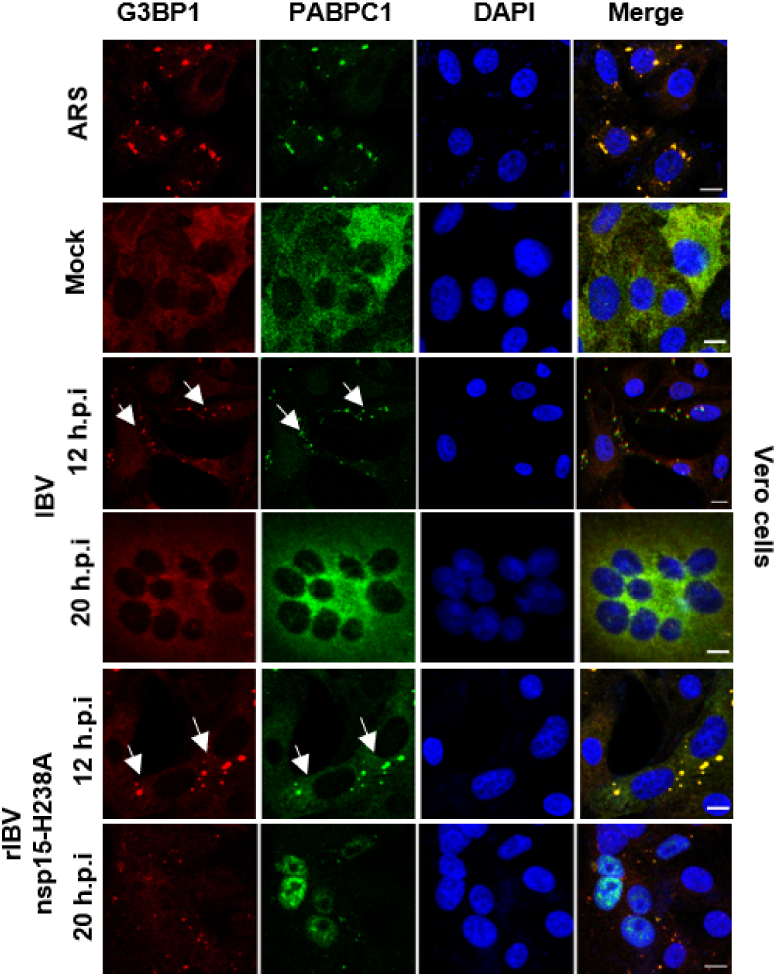
Infection with rIBV-nsp15-H238A, but not with IBV-WT, triggers PABPC1 nuclear retention. In the infected cells with SG formation, PABPC1 aggregates to SG. (A) Vero cells were infected with IBV-WT or rIBV-nsp15-H1238A at an MOI of 1. At 12 and 20 h.p.i, indirect immunofluorescence was performed to detect G3BP1 (red), PABPC1 (green) and nuclei were stained with DAPI (blue). White arrows indicate the infected cells with SG formation and SG localization of PABPC1.

**Supplementary figure 5.**
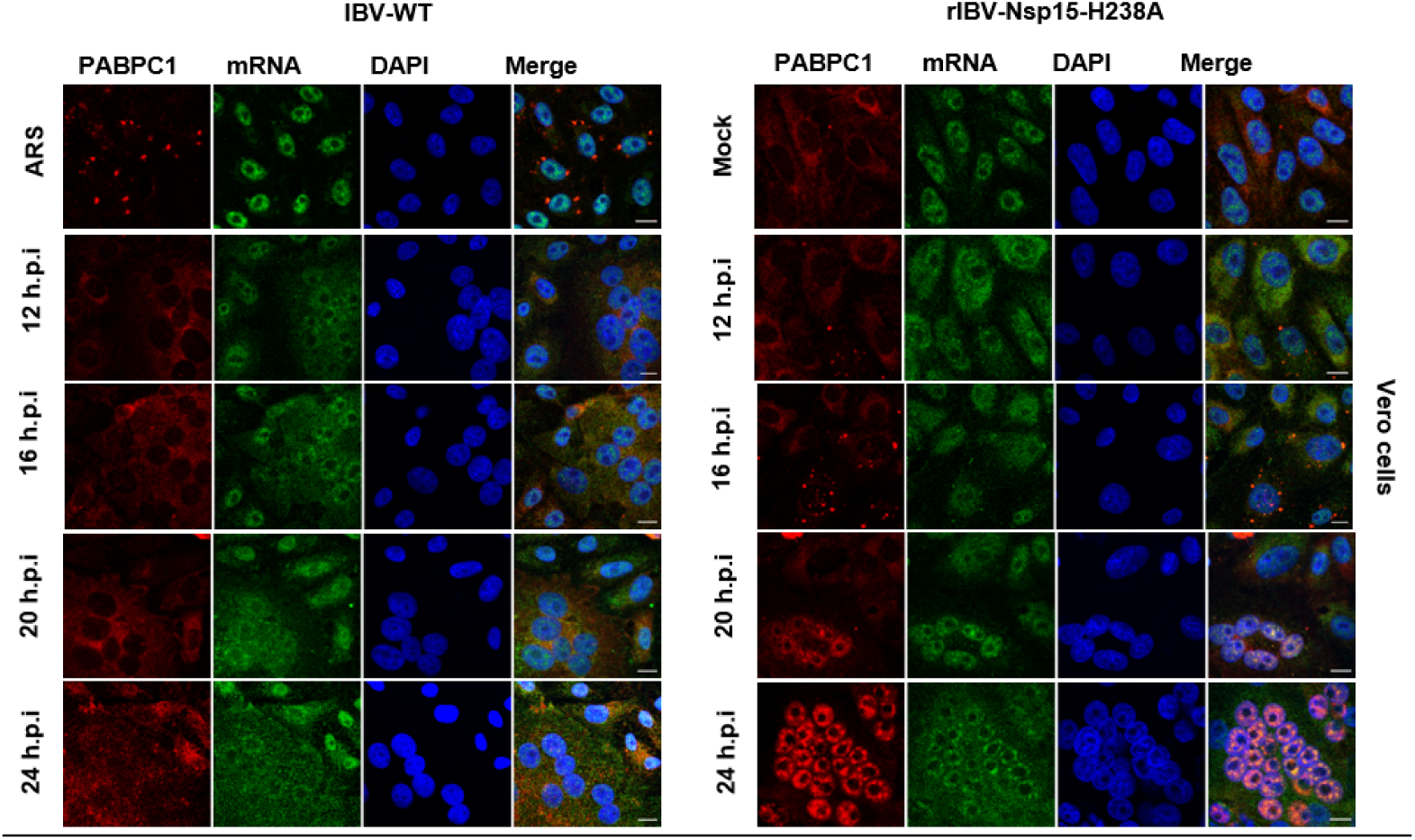
Infection with rIBV-nsp15-H238A, but not with IBV-WT, triggers nuclear retention or SG localization of both PABPC1 and mRNA. (A) Vero cells were infected with IBV-WT (left panel) or rIBV-nsp15-H1238A (right panel) at an MOI of 1. At 12, 16, 20 and 24 h.p.i, FISH and indirect immunofluorescence were performed to detected PABPC1(red) and mRNA (green). Nuclei were stained with DAPI (blue).

## Notes

### Competing Interest Statement

The authors have declared no competing interest.

